# Transcriptomic and functional analysis of Aβ_1-42_ oligomer-stimulated human monocyte-derived microglia-like cells

**DOI:** 10.1101/2021.08.12.456055

**Authors:** Tamar Smit, Paul R. Ormel, Jacqueline A. Sluijs, Lianne A. Hulshof, Jinte Middeldorp, Lot D. de Witte, Elly M. Hol, Vanessa Donega

## Abstract

Dysregulation of microglial function contributes to Alzheimer’s disease (AD) pathogenesis. Several genetic and transcriptome studies have revealed microglia specific genetic risk factors, and changes in microglia expression profiles in AD pathogenesis, *viz*. the human-Alzheimer’s microglia/myeloid (HAM) profile in AD patients and the disease-associated microglia profile (DAM) in AD mouse models. The transcriptional changes involve genes in immune and inflammatory pathways, and in pathways associated with Aβ clearance. Aβ oligomers have been suggested to be the initial trigger of microglia activation in AD. To study the direct response to Aβ oligomers exposure, we assessed changes in gene expression in an *in vitro* model for microglia, the human monocyte-derived microglial-like (MDMi) cells. We confirmed the initiation of an inflammatory profile following LPS stimulation, based on increased expression of *IL1B, IL6*, and *TNFα*. In contrast, the Aβ_1-42_ oligomers did not induce an inflammatory profile or a classical HAM or DAM profile. Interestingly, we observed a specific increase in the expression of metallothioneins in the Aβ_1-42_ oligomer treated MDMi cells. Metallothioneins are involved in metal ion regulation, protection against reactive oxygen species, and have anti-inflammatory properties. In conclusion, our data suggests that Aβ_1-42_ oligomers may trigger a protective response both *in vitro* and *in vivo*.

## Introduction

Alzheimer’s disease (AD) is a neurodegenerative disorder that is clinically characterized by progressive memory loss and impairment in cognitive functions (Querfurth and LaFerla, 2010). Hallmarks of AD pathology are the aggregation of amyloid-beta (Aβ) in extracellular plaques, intraneuronal hyperphosphorylated tau tangles (Hardy and Allsop, 1991; Selkoe, 1991), and reactive gliosis (Itagaki et al., 1989; Kato et al., 1998). Aβ plaques are surrounded and infiltrated by reactive microglia (Itagaki et al., 1989; Rozemuller et al., 1986).

Microglia are the resident immune cells of the brain (Ginhoux et al., 2013, 2010). In neurodegenerative diseases, including AD, microglia adopt an activated phenotype and undergo major morphological and functional changes (Wolf et al., 2017). Genome-wide association studies identified several AD risk loci that are found in or near genes predominantly expressed in microglia (*APOE, TREM2*, and *CD33*) (Hemonnot et al., 2019; Johnson et al., 2020; Lambert et al., 2013). The identification of a distinct AD-pathology-associated gene expression profile in microglia by recent transcriptome studies support the important role of microglia in AD (Mathys et al., 2019; Srinivasan et al., 2019). The AD-pathology-associated microglia profile was enriched in immune and inflammatory pathways, as well as in Aβ clearance pathways (Mathys et al., 2019). Srinivasan *et al*., 2019 referred to their Human Alzheimer’s Microglia/Myeloid cells profile as the HAM signature (Srinivasan et al., 2019). Single-cell RNA sequencing analysis of sorted microglia from the 5xFAD AD mouse model revealed a distinct microglia subtype, referred to as disease-associated microglia (DAM) (Keren-Shaul et al., 2017). Some well-known AD risk factors, including *ApoE, Ctsd, Lpl, TYROBP*, and *TREM2* are upregulated in DAM (Keren-Shaul et al., 2017).

In AD, the acute activation of microglia by Aβ can be beneficial, as it increases phagocytosis and clearance of Aβ by microglia (Prokop et al., 2013; Rogers et al., 2002). However, when the activation of microglia becomes chronic, the cells contribute to neurotoxicity by the release of pro-inflammatory cytokines and mediate synapse loss (Bamberger et al., 2003; Sarlus and Heneka, 2017). Aβ in the form of monomers, oligomers, and fibrils, has multiple biological effects, including neurotoxicity and the activation of microglia (Lue et al., 2019; Walker et al., 2006, 2001). The Aβ oligomers are considered to be the main culprit in AD pathogenesis events (Esparza et al., 2013; Selkoe and Hardy, 2016).

In this study, we investigated the direct effect of Aβ_1-42_ oligomers on the microglia transcriptome. We decided not to use microglia from aged human brain tissue as these are likely to already have been exposed to Aβ. Furthermore, the isolation procedure of microglia from *post-mortem* human brain tissue changes its activation state, leading to the loss of classical microglial marker expression once cultured (Gosselin et al., 2017). Therefore, we used the human monocyte-derived induced microglia-like (MDMi) cell model as a source of human microglia to investigate the transcriptomic changes induced by human Aβ_1-42_ oligomer stimulation. The MDMi cell model is based on established protocols to differentiate human monocytes towards a microglia-like phenotype (Leone et al., 2006; Ohgidani et al., 2015; Ryan et al., 2017), with minor adaptations (Ormel et al., 2020). In short, monocytes isolated from healthy controls are differentiated in MDMi cells using several factors that are important for microglia development. Studies have shown that the MDMi cell models display characteristics of central nervous system resident microglia and can be used for transcriptome analysis and functional assays (Leone et al., 2006; Ohgidani et al., 2015; Ryan et al., 2017; Sellgren et al., 2019). These findings confirm that MDMi cells can be used as a model to study human microglia function in health and disease.

We first describe the generation of the MDMi cell model and investigate their response to the classical inflammatory stimulus lipopolysaccharide (LPS), which confirmed the initiation of an inflammatory profile, based on an increased expression of *IL1B, IL6*, and *TNFα*. We then exposed the MDMi cells to Aβ_1-42_ oligomers to study changes in the transcriptome and determined whether the expression of the most significant upregulated genes was also increased in human primary microglia isolated from AD compared to non-demented control (NDC) cases. We also compared three well-known AD microglia gene profiles (Keren-Shaul et al., 2017; Mathys et al., 2019; Srinivasan et al., 2019) to our stimulated and unstimulated MDMi cells. Our results showed that the Aβ_1-42_ oligomers did not induce an inflammatory profile or a classical HAM or DAM profile. Instead, we observed a specific increase in the expression of metallothionein genes following Aβ_1-42_ oligomer treatment. Our work suggests that Aβ_1-42_ oligomers may initially induce a protective response in MDMi cells.

## Materials and methods

### Data and code availability

The bulk RNA sequencing dataset generated in this study have been deposited in NCBI’s Gene Expression Omnibus (accession number:). All the analysis is described in the Methods.

### Differentiation of monocyte-derived microglia-like (MDMi) cells

#### Monocyte isolation

Blood samples of 10 healthy controls were obtained from the Dutch blood bank (Sanquin, https://www.sanquin.nl/en) (Supplementary Table 1). Peripheral blood mononuclear cells (PBMCs) were enriched by density gradient separation using Ficoll (Ficoll-Paque™ plus, GE Healthcare). Monocytes were isolated from PBMCs with anti-CD14 conjugated magnetic microbeads (130-050-201, Miltenyi Biotec) according to the manufacturer’s protocol. Monocytes were immersed in 45% fetal calf serum (FCS, 10500 ThermoFisher Scientific), and 10% dimethyl sulfoxide (DMSO) in RPMI culture medium (RPMI 1640, Gibco with 100 Units/mL penicillin, 100 μg/mL streptomycin and 2mM L-glutamine) and stored in liquid nitrogen until further use.

#### Monocyte-derived microglia-like cell establishment

The protocol for generating MDMi cells is based on established protocols (Leone et al., 2006; Ohgidani et al., 2015; Ryan et al., 2017), with minor adaptations to optimize the expression of *TREM2, TYROBP* and *PROS1* (Ormel et al., 2020). To generate MDMi cells, monocytes were thawed on ice-cold RPMI medium and plated at a density of 600.000 cells/well or 200.000 cells/well in a 48- or 96-well plate for transcriptome and phagocytosis assays, respectively. The wells were coated with poly-L lysine (PLL, Sigma Aldrich) at 37 °C in 5% CO_2_ for 30 minutes. Cells were left to adhere to the wells under standard humified culture conditions at 37 °C in 5% CO_2_ for at least 1 hr, then cells were washed with PBS (Invitrogen) and the medium was replaced by 25% ACM (Astrocyte conditioned medium SCC1811, Sanbio) in RPMI medium. To induce the differentiation towards MDMi cells, the 25% ACM/RPMI medium was replaced on day four and day eight with RPMI containing 25% ACM, 10 ng/ml human M-CSF (130-096-491, Miltenyi Biotec), 10 ng/mL GM-CSF (130-093-862, Miltenyi Biotec), 1 ng/ml TGFβ (130-095-067, Miltenyi Biotec), 12.5 ng/ml IFN-γ (130-096-872, Miltenyi Biotec), and 100 ng/mL IL-34 (130-108-997, Miltenyi Biotec).

#### Stimulation of MDMi cells with Aβ_1-42_ oligomers or lipopolysaccharide

At day 10 in culture, MDMi cells were treated with 500 nM stable human Aβ_1-42_ oligomers (180222EMH, gift from Crossbeta) or 100 ng/mL lipopolysaccharide in PBS (LPS, L4391-1MG, E. coli 0111:B4, Sigma-Aldrich). As a control for the oligomer and LPS condition, the same volume of vehicle (20 mM HEPES, 150 mM NaCl and 200 mM sucrose, pH 7.2) or PBS was added, respectively. After 24 hrs of stimulation, MDMi cells were washed with PBS and collected for RNA isolation (48-well plates) in Trizol (Ambion, Life Technologies, Carlsbad, CA, USA) or used for phagocytosis assay.

### RNA isolation, library preparation, and sequencing

RNA isolation was performed using the miRNeasy Mini kit (217004, Qiagen) according to the manufacturer’s protocol including the DNase treatment. The RNA concentration was determined using a Nanodrop 2000 spectrophotometer (Thermo Scientific, Waltham). cDNA libraries were prepared according to the CelSeq2 protocol (Hashimshony et al., 2012; Simmini et al., 2014) by Single Cell Discoveries, Utrecht (Muraro et al., 2016). Briefly, for each sample a custom-made primer was added to the RNA, denatured at 70 °C for 2 min, and immediately cooled down. mRNA was reverse transcribed into cDNA using clean-up beads (ThermoFisher Scientific, Ambion, Waltham, USA). The purified cDNA was transcribed *in vitro* to obtain amplified RNA using the MegaScript kit, which was followed by purification with Agencourt RNAclean XP (RNAse free) beads (Beckman Coulter). Amplified RNA was fragmented with fragmentation buffer and purified using the RNA cleanXP beads. Quality of the amplified RNA was determined with the RNA 6000 Pico chips (Agilent) on an Agilent 2100 bioanalyzer. Next, the amplified RNA was reverse transcribed and amplified by PCR. Libraries were labelled with a 4 bp unique molecular identifier (UMI) that was added to the primer. Samples from the different stimulation and control conditions were pooled in libraries for each donor (see Supplementary Table 1). The quality of the libraries was determined with the DNA High Sensitivity chips on an Agilent 2100 bioanalyzer (Agilent). Libraries were sequenced on the Illumina NextSeq 500 platform using paired-end sequencing (75 bp) with a depth of 10M reads per sample.

### Bulk RNA sequencing analysis

Libraries were de-multiplexed and raw reads were aligned to the hg19 human RefSeq transcriptome with Burrows-Wheeler Aligner (BWA) (Li and Durbin, 2010). Duplicate reads and reads that mapped equally well to multiple locations were discarded. The quality control, normalization, and identification of differentially expressed genes were done with DESeq2, an R (version 3.6.3) based package (Love MI et al 2014). Samples that had more than 10M reads were discarded (see Supplementary Table 1). Read counts were normalized to transcripts per million (TPM). The gene expression was corrected for the covariate sex. Genes were considered differentially expressed with an adjusted (adj.) p-value of less than 0.05 and a Log2 fold change of at least 2 for the comparison LPS and PBS and a Log2 fold change of at least 0.5 for Aβ_1-42_ oligomers and vehicle stimulation.

### Gene ontology enrichment and heatmap analysis

The gene ontology (GO) (GO Molecular Function 2018) and Panther pathway analyses (Panther 2016) were performed on the list of differentially expressed genes. The lists of genes used for analysis are provided as Supplementary Table 2 and 3. We loaded these genes in EnrichR (http://amp.pharm.mssm.edu/Enrichr/), a web-based tool for enrichment analysis (Chen et al., 2013; Kuleshov et al., 2016). Heatmaps were created using the Morpheus Broad Institute Software (https://software.broadinstitute.org/morpheus/).

### Comparison with previously published datasets

*Srinivasan et al., 2019 – Human Alzheimer’s microglia/myeloid cells (HAM) profile* - Differential gene expression results of myeloid cells of AD and control subjects were downloaded from http://research-pub.gene.com/BrainMyeloidLandscape/BrainMyeloidLandscape2/#study/study/GSE125050/studyReport.html (see Supplementary Table 4). Briefly, Srinivasan *et al*., 2019 performed RNA sequencing on FAC-sorted cells from frozen *post-mortem* brain tissue. Cells were isolated from fusiform gyrus tissue from 10 AD and 15 control cases (Srinivasan et al., 2019).

*Mathys et al., 2019 –* Differential gene expression results of microglia were obtained from Supplementary Table 2 and 7 (Mathys et al., 2019). Mathys *et al*., 2019 performed single-nuclear RNA sequencing on nuclei from frozen *post-mortem* tissue of 24 AD and 24 no-pathology cases. A total of 1920 microglia nuclei were obtained (Mathys et al., 2019).

*Keren-Shaul et al., 2017 – Disease-associated microglia (DAM) profile* - The differential gene expression data of isolated cortical cells from three 5xFAD transgenic and three wild-type 6-month-old mice were obtained from Supplementary Table 2 (Keren-Shaul et al., 2017). This study identified 500 differentially expressed genes that make-up the disease-associated microglia (DAM) profile with single-cell RNA sequencing (Supplementary Table 4) (Keren-Shaul et al., 2017).

### Phagocytosis assay

Fluosphere carboxylate-modified microspheres (2 μm, yellow-green fluorescent, F8827, ThermoFisher) were used for the phagocytosis assay. After 10 days of differentiation, MDMi cells were stimulated with Aβ_1-42_ oligomers or LPS for 24 hrs. In the unstimulated control conditions vehicle (20 mM HEPES, 150 mM NaCl and 200 mM sucrose, pH 7.2) or PBS was added to the culture medium. Next, the MDMi cells were cultured with the uncoated fluorescent beads (3 beads/cell) under standard humified culture conditions at 37 °C in 5% CO_2_ for 1 hr. For each condition, three cell culture wells were used per donor. After 1 hr incubation with the beads, the cells were washed twice with cold PBS to remove non-phagocytized beads. The cells were detached using Trypsin 1x (15090, ThermoFisher Scientific) and 0.5 M EDTA, pH 8.0 in PBS under standard humified culture conditions at 37 °C in 5% CO_2_ for 5 min. Fluorescence activated cell sorting (FACS, BD FACSCanto™ II, software version 8.0.1) was used to analyze >1000 living cells, gated by FSC and SSC. Phagocytosis of fluorescent beads was gated by FSC and the blue 1-a (laser 488, filter 530/30) channel. The gates were set by first sorting MDMi cells without beads, monocytes, and beads only. These gate settings were later used to sort the positive MDMi cells.

### Immunocytochemistry

Monocytes were plated on PLL-coated coverslips in a 24-well plate. After 10 days of differentiation, MDMi cells were washed with PBS containing 137 mM NaCl, 1.8 mM KH_2_PO_4_, 5.96 mM Na_2_HPO_4_.2H_2_O, 2.7 mM KCl, pH 7.4 and fixed in 4% PFA. MDMi cells were incubated with a primary antibody against Iba1 (1:1,000; 019-19744, FUJIFILM Wako Chemicals) in blocking buffer (2% bovine serum albumin, 0.1% TritonX100 (Merck, Darmstadt, Germany), and 5% normal donkey serum (017-000-121, Jackson ImmunoResearch) in PBS). Following washes with PBS, the secondary antibody donkey-anti-rabbit 488 (1:700; 715-585-150, Jackson ImmunoResearch) and Hoechst (1:1,000; H3569, ThermoFischer Scientific) as a nuclear staining were added. Finally, MDMi cells were washed with PBS and embedded with Mowiol (0.1 M Tris-HCl, pH 8.5, 25% Glycerol, 10% w/v Mowiol 4-88). Images were taken on a Zeiss AxioScopeA1 microscope using an EC Plan-Neofluar 20x/0.50 M27 objective, an AxioCam MRm camera (Zeiss), and Zen 2011 software. Images were taken with a resolution of 1388 × 1040 pixels. The phagocytosis assay was imaged with a Zeiss LSM 880 confocal laser microscope using a 40x/1.3NA oil DICII objective (EC PlnN), and Zen black Z.1SP3 software. Images were taken with a z-step of 0.7 μm and a resolution of 1024 × 1024 pixels. The phagocytosis assay was imaged with a Zeiss LSM 880 confocal laser microscope using a 40x/1.3NA oil DICII objective (EC PlnN), and Zen black Z.1SP3 software. Images were taken with a z-step of 0.7 μm and a resolution of 1024 × 1024 pixels.

### Isolation of human microglia

Human brain tissue was obtained from the Netherlands Brain Bank (NBB, www.brainbank.nl). Permission for brain autopsy and the use of brain tissue and clinical information for research purposes were obtained per donor *ante-mortem* by the NBB. The donor identity was pseudo-anonymized by the NBB. For some donors, the amyloid and Braak scores still have to be confirmed by a neuropathologist (Supplementary Table 5).

Our microglia isolation protocol was adapted from Melief *et al*., 2016. Briefly, the isolation of microglia started 6 to 24 hrs after autopsy. Approximately 4-10 grams of tissue of the gyrus temporalis superior (GTS1-3) was collected in cold Hibernate-A medium. GTS1-3 tissue was mechanically dissociated with a scalpel in ice-cold GKN-BSA containing 11.1 mM D-(+)-Glucose monohydrate (14431-43-7, Sigma Aldrich) and 50 mM bovine serum albumin (BSA, A450-3, Sigma-Aldrich) in PBS, pH 7.4 (10010031, Invitrogen). To obtain a cell suspension, tissue was enzymatically dissociated with Collagenase type I (370 units/ml, LS004196, Worthington Biochemical) and DNase I (100 μg/ml, 10104159001, Sigma-Aldrich) at 37 °C in a shaking incubator (140-170 rpm) for 1 hr. Dissociated GTS tissue was washed and centrifuged at 1,800 rpm at 4 °C, and the pellet was resuspended in GKN-BSA. The cell suspension was filtered through a 100 μm cell strainer (Corning, New York, USA). Percoll (GE Healthcare) gradient centrifugation was used to separate the different cellular fractions. Percoll was added dropwise to the cells in GKN-BSA and centrifuged at 4000 rpm at 4 °C for 30 min. The middle turbid layer (cellular fraction) was carefully collected and washed with an equal volume of GKN-BSA, centrifuged and suspended in MACS buffer containing 2 mM EDTA and 1% FBS in PBS 1x (Invitrogen). Microglia were isolated with anti-CD11b conjugated magnetic microbeads (130-049-601, Miltenyi Biotec GmbH) according to manufacturer’s protocol, using MS columns (130-042-201, Miltenyi Biotec) placed in a magnetic field. The eluted CD11b positive cells were stored in Trizol until RNA isolation for qPCR.

### Isolation of mouse microglia

All experiments were performed in line with institutional guidelines of the University Medical Center Utrecht, approved by the Animal Ethics Committee of Utrecht University (AVD1150020174314), and were conducted in agreement with Dutch laws (wet op de Dierproeven, 1996) and European regulations (Guidelines 86/609/EEC). Animals were housed under standard conditions with access to water and food *ad libitum*. We used the well-established AD mouse model, APPswePS1dE9 double transgenic line (Jankowsky et al., 2001; Orre et al., 2014). This line has been backcrossed to C57BL/6 mice for more than 20 generations (Kamphuis et al., 2015), since the genetic background is known to influence AD pathogenesis (Hyman and Tanzi, 2019; Song et al., 2011; Tahara et al., 2006). We isolated microglia from three pooled cortices of 4-month-old AD mice and wild-type littermates as controls (resulting in N = 12 samples). Genotype was confirmed by performing real-time PCR with primers targeted to the two transgenes expressed by the APP/PS1 mice — human/mouse chimeric APP with K595N/M596L Swedish mutation and human PS1 carrying the Exon 9 deletion. For further details on this transgenic line see The Jackson Laboratory (B6C3-Tg(APPswe,PSEN1dE9)85Dbo/Mmjax;StockNo: 34829; https://www.jax.org/strain/004462). Mice were anesthetized by an overdose of Pentobarbital and transcardially perfused with HBSS (14175-053, Gibco). Cortical regions were dissected on ice and immediately processed for transcriptome analysis. Briefly, cortical tissue was mechanically dissociated and subjected to enzymatic dissociation using Papain (final concentration of 8 U/ml, Worthington) in combination with 100 μg/ml DNase I in Pipes based buffer containing 1 mM Pipes (P1851, Sigma Aldrich), 25 mM L-Cystein HCL, and 5 mM EDTA. Tissue was enzymatically dissociated at 37 °C in a shaking incubator (Incu-shaker mini, Benchmark) for 50 min. Subsequently, DNase I in GKN/BSA was added and the tissue suspension was incubated for another 15 min. A 90% Percoll gradient centrifugation was used to collect the cellular fraction before cells were incubated with anti-CD11b conjugated magnetic microbeads according to the manufacturer’s protocol. The eluted CD11b positive cells were stored in Trizol until further use.

### RNA isolation, cDNA synthesis, and quantitative real-time PCR for human and mouse microglia

For RNA isolation, samples were thawed and total RNA was isolated with TRIzol (Life Technologies) according to manufacturer’s protocol, and the RNA was subsequently precipitated in 2-propanol and 20 μg/μl glycogen (Roche) overnight at −20 °C. Samples were centrifuged (12,000 x g for 30 minutes) at 4 °C, washed twice with cold 75% ethanol, and the RNA pellet was dissolved in MilliQ. The RNA concentration was determined using a Nanodrop 2000 spectrophotometer (Thermo Scientific, Waltham, MA, USA).

Total RNA was treated with DNaseI (gDNA wipe out buffer, Qiagen) at 42 °C for 2 min, and cDNA was synthesized using Quantic Reverse Transcription kit (Qiagen, Hilden, Germany) following the manufacturer’s instructions. Quantitative PCR (qPCR) was performed using the Quantstudio 6 Flex (Applied Biosystems, Life Technologies, USA). For the qPCR reaction 1 μl of 1:20 diluted cDNA in MilliQ was used with 1 μl primer mix (forward and reverse primers, 2 pmol/μl, Supplementary Table 7), 5 μl FastStart Universal SYBR Green Master mix (Roche, Basel, Switzerland) and 3 μl MilliQ. The following cycling conditions were used: 2 min 50 °C, 10 min 95 °C, and 40 cycles of 15 sec 95 °C and 1 min 60 °C. A dissociation curve was obtained by ramping the temperature from 60 °C to 90 °C. Applied Biosystems 7500 Real-Time PCR software (Applied Biosystems) was used for the analysis of amplification curves. Gene expression was normalized to three of the following reference genes *β-actin, S18*, hypoxanthine phosphoribosyltransferase (*HPRT*) and glyceraldehyde-3-phosphate dehydrogenase (*GAPDH*).

### Statistics

Data was analyzed with GraphPad Prism8. A paired two-sided t-test was used for groups with equal variances or its nonparametric equivalent, the Wilcoxon signed rank test was used to determine the effect of the different stimulations on MDMi cells. For the statistical analysis of mouse microglia and human microglia data, the unpaired two-sided t-test for two-group comparison for groups with equal variances was used. The one-way ANOVA followed by Bonferroni’s *post-hoc* test was used to assess the effect between three groups. Normal distribution was tested with the D’Agostino & Pearson omnibus normality test. For data that were not normally distributed the Mann-Whitney test or Kruskal-Wallis test was used followed by Dunn’s tests for multiple comparisons. Outliers were identified using the Robust regression and Outlier removal (ROUT) method with the coefficient set to 1, available in GraphPad.

## Results

### Characterization of the monocyte-derived microglia-like cell (MDMi) model

Human monocytes from 10 healthy individuals were cultured for 10 days and differentiated into a microglia-like phenotype with M-CSF, GM-CSF, IL-34, TGFβ, and IFN-γ, which are important factors in the development of microglia (Ryan et al., 2017). This led to the development of a microglia-like morphology, *i*.*e*. a ramified appearance and the expression of Iba1 (Fig. 1a-b). These monocyte-derived microglia-like (MDMi) cells had a distinct gene expression profile compared to monocytes based on increased expression of the common microglial markers, *APOE, TREM2*, and *C1QA* (Fig. 1c). Other microglial markers, such as *P2YR12*, were not significantly upregulated in MDMi cells when compared to monocytes. We also compared the gene expression profile of primary human microglia (pMG) to MDMi cells. Expression levels of *IRF8, TREM2*, and *C1QA* were similar in MDMi and pMG. These findings are in line with previous work, showing that unsupervised clustering based on the expression of a microglial marker panel resulted in clustering of cultured pMG with MDMi cells and less with monocytes (Ormel et al., 2020). Based on both morphology and gene expression, MDMi cells resemble primary human microglia and therefore can be used as a model to study human microglia function in health and disease. To characterize the MDMi cells further, we determined the phagocytic function of MDMi cells and compared that to pMG. Uncoated fluorescent beads were taken up by MDMi cells after 1 hr of incubation with the beads (Fig. 1d). Fluorescence activated cell sorting (FACS) was used to determine the percentage of phagocytizing cells, *i*.*e*. the phagocytosis index (Fig. 1e). Our results show that MDMi cells, like microglia, can phagocytize, although significantly less than pMG (24.5% ± 5.5 vs 10.3% ± 1.1; Fig. 1f).

**Figure 1.**
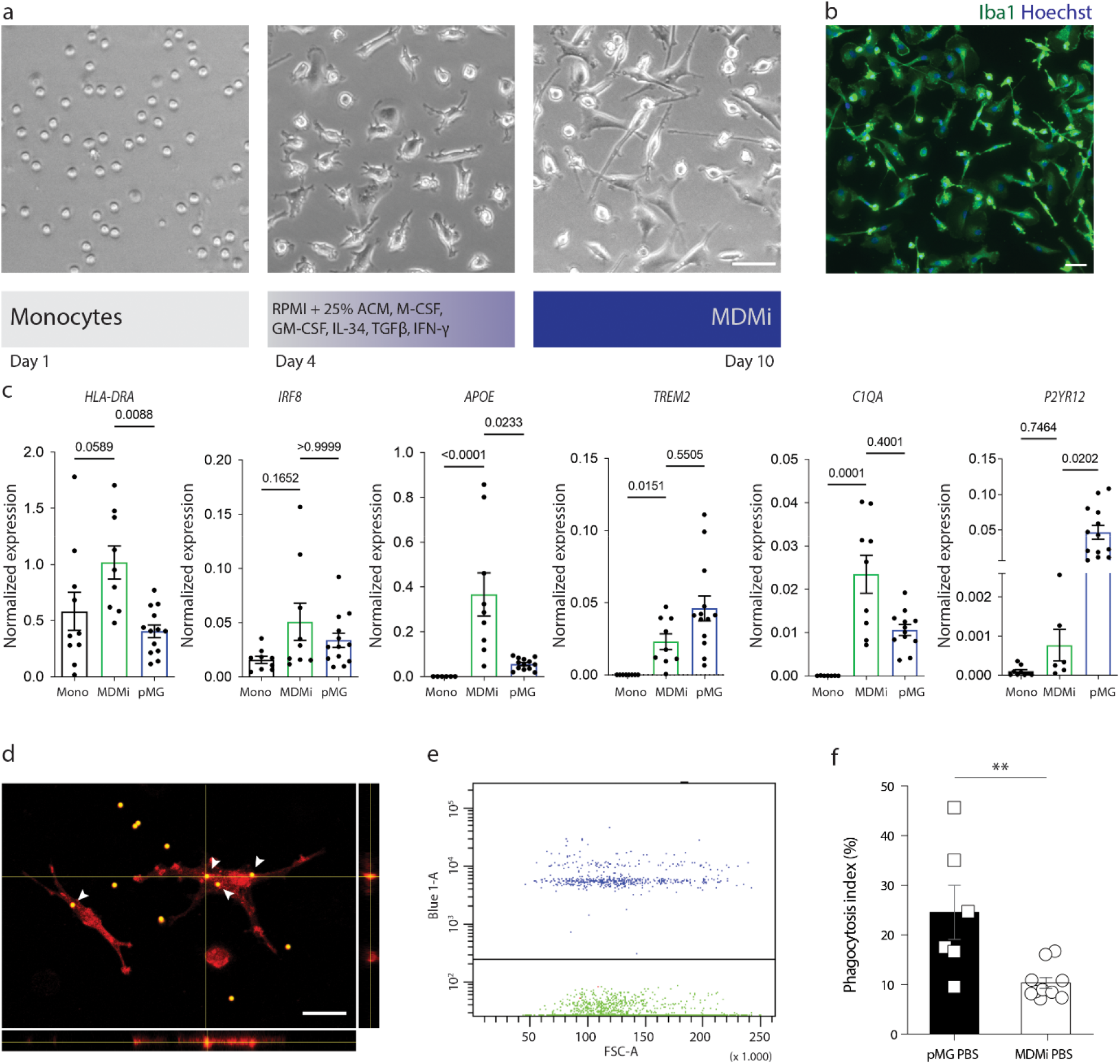
Characterization of monocyte-derived microglia-like cells. a. Representative bright-field images of the differentiation of monocytes into monocyte-derived microglia-like (MDMi) cells and scheme of the MDMi differentiation protocol. Cells were cultured for 10 days with differentiation medium, containing M-CSF, GM-CSF, IL-34, TGF-β, and IFN-γ. Scale bar is 40 μm. b. Iba1 immunostaining together with the nuclear staining (Hoechst) of MDMi cells 10 days in culture. Scale bar is 40 μm. c. mRNA expression of common microglial markers in monocytes (Mono), MDMi cells, and primary human microglia (pMG). d. MDMi cells were cultured in the presence of fluorescent beads (2 μm) for 1 hour. Confocal image of MDMi cells (Iba1, red) with inclusion of fluorescent beads (yellow). Arrowheads show beads that were taken up by MDMi cells. Scale bar is 40 μm. e. A representative plot of the flow cytometry gates used to quantify the phagocytosis of fluorescent beads by MDMi cells after 24 hrs exposure to PBS. Cells were plotted on a forward scatter and Blue1-A plot to visualize the cells with inclusion of one or more fluorescent beads (blue dots) and MDMi cells without internalized beads (green dots). f. Quantification of phagocytosis by pMG and MDMi cells (t-test, *p* = 0.006). Data represented as mean ± SEM.

### Inflammatory response of monocyte-derived microglia-like cells

To assess whether LPS stimulation induced changes in the transcriptome we performed bulk RNA sequencing on stimulated and unstimulated MDMi cells. We first confirmed that MDMi cells expressed genes that encode for proteins that are essential for an LPS-induced immune response, such as the toll-like receptor-4 (*TLR4*), *CD14*, and *Ly96* (Fig. 2a). Next, we used the phagocytosis assay to assess the effect of LPS stimulation on the phagocytic ability of MDMi cells. The 24 hrs stimulation with 100 ng/ml LPS increased the phagocytosis index of the MDMi cells by 7% (PBS 10.3% ± 1.1; LPS 17.9% ± 1.8; Fig. 2b). Whole transcriptome analysis showed an inflammatory response in MDMi cells after LPS stimulation. A principal component analysis (PCA) identified two clusters corresponding to the LPS stimulated and unstimulated MDMi samples (PBS; Fig. 2c). Differential gene expression analysis identified 675 genes that were significantly upregulated (Log2 FC > 2 and adj. *p*-value < 0.05) and 413 genes that were downregulated (Log2 FC > −2 and adj. *p*-value < 0.05) after LPS stimulation (Supplementary Table 2).

**Figure 2.**
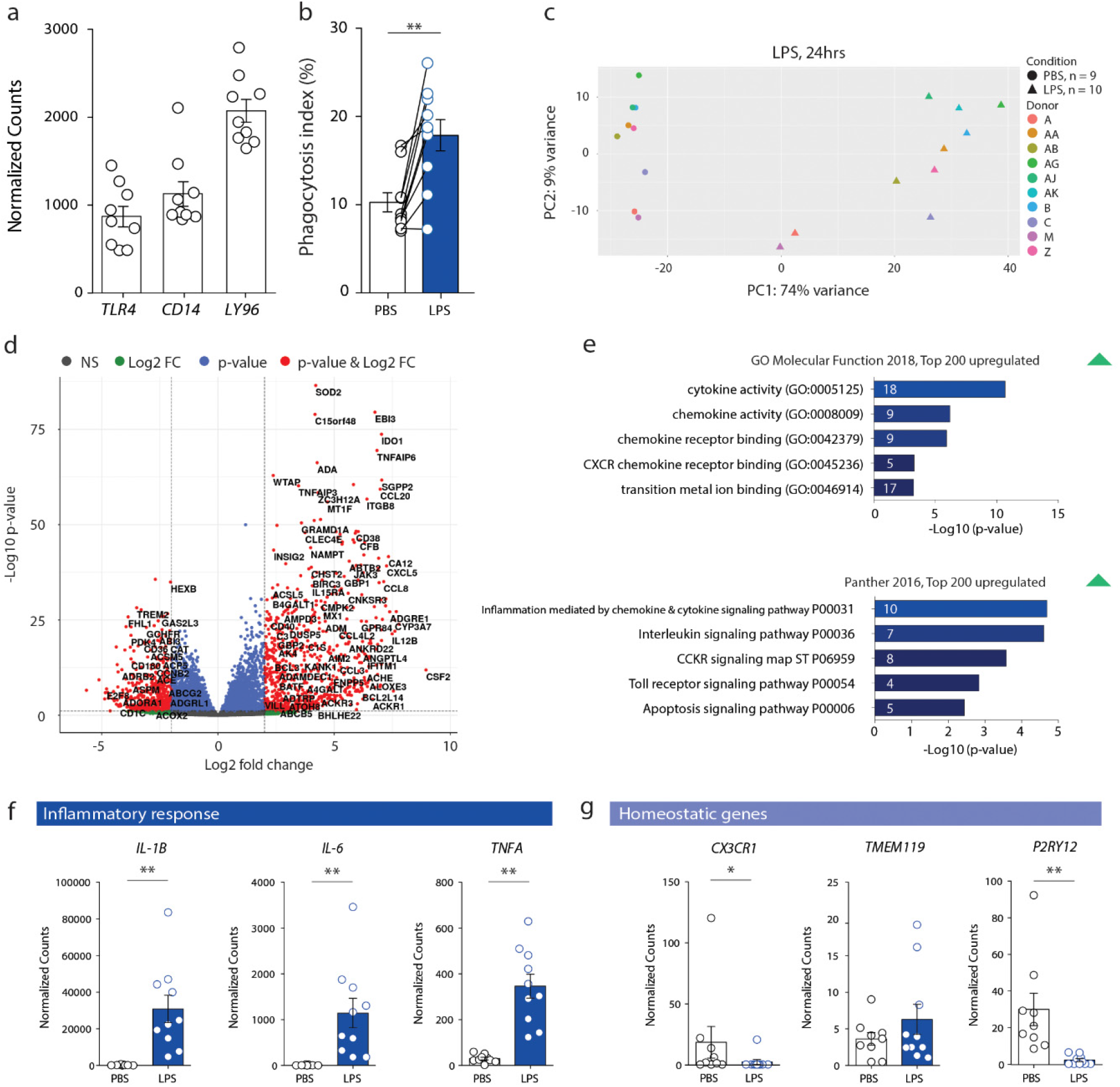
LPS response in monocyte-derived microglia-like cells. MDMi cells of 10 donors were stimulated for 24 hrs with 100 ng/ml lipopolysaccharide (LPS) and used for RNA sequencing. a. TPM-normalized read counts of genes, encoding for proteins involved in LPS binding, determined in unstimulated MDMi cells (PBS condition). b. Quantification of phagocytosis index (paired t-test, *p* = 0.001). Each dot in the graphs represents the mean of three cell culture wells/donor. Data are presented as mean ± SEM. c. Principal component analysis of paired unstimulated and LPS stimulated MDMi samples. d. Volcano plot showing differential gene expression after LPS stimulation of the MDMi cells (genes with Log2 fold change > 2 and *p* < 0.05 shown in red). e. Gene ontology and pathway analyses performed on the top 200 most significantly upregulated genes (Log2 fold change of 2 and adj. *p* < 0.05) in MDMi cells after LPS stimulation. The number of genes that are associated with each GO term are shown within the bars. f. TPM-normalized read counts of a selection of inflammatory response genes (*IL1B, IL6*, and *TNFα*) expressed by MDMi cells that are significantly upregulated by LPS stimulation. g. TPM-normalized read counts of a selection of homeostatic genes (*CX3CR1, TMEM119*, and *P2RY12*) expressed in MDMi cells.

Gene ontology (GO) and Panther pathway analyses on the 200 most significantly upregulated differentially expressed genes after LPS stimulation, identified enrichment for genes involved in cytokine, chemokine, and interleukin signaling pathways (Fig. 2e). The genes enriched in these pathways, such as *IL1B, IL6, TNFα* (Fig. 2f), and several chemokines are known to be upregulated in primary microglia after stimulation by LPS (Melief et al., 2016; Sneeboer et al., 2019). Moreover, the GO analysis also revealed changes in “transition metal ion binding” due to upregulation of the expression of *SOD2, IL1α*, and two isoforms of the metallothionein family, *MT1* and *MT2*.

The expression of some microglia homeostatic markers, including *CX3CR1* and *P2RY12*, were downregulated by LPS stimulation, while *TMEM119* remained unaffected (Fig. 2g). This is in line with previous studies showing a downregulation of homeostatic microglial genes after stimulation (Butovsky and Weiner, 2018; Keren-Shaul et al., 2017). Overall, LPS stimulation of MDMi cells resulted in an immune activated phenotype.

### Transcriptomic changes induced by Aβ_1-42_ oligomer stimulation

We performed bulk RNA sequencing on stimulated and unstimulated MDMi cells to assess the direct effect of Aβ_1-42_ oligomer stimulation on changes in the transcriptome. We confirmed that the MDMi cells express several genes, including *CD33, TLRs*, formyl peptide receptor 2 (*FPR2*), and *TREM2*, encoding for proteins involved in Aβ clearance and in triggering an inflammatory response (Fig. 3a) (Doens and Fernández, 2014; Salminen et al., 2009; Tejera and Heneka, 2016). In line with this, stimulating the MDMi cells with 500 nM Aβ_1-42_ oligomers for 24 hrs increased the phagocytosis index of the MDMi cells with 5.7% (Vehicle 9.8% ± 1.4; Oligomers 15.5% ± 1.9; Fig. 3b).

**Figure 3.**
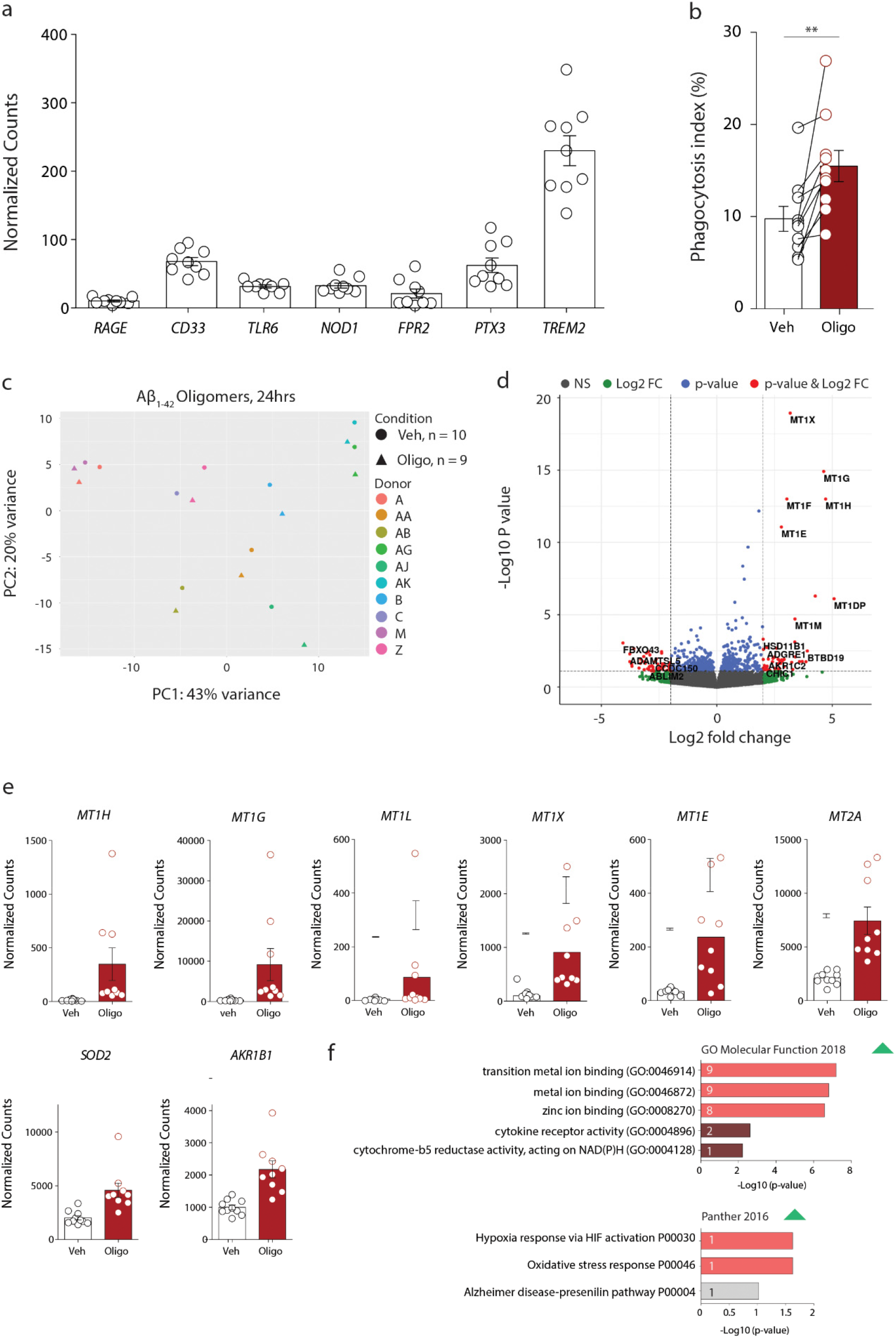
Aβ_1-42_ oligomer stimulation of monocyte-derived microglia-like cells. a. TPM-normalized read counts in unstimulated MDMi cells of genes encoding common receptors for Aβ. b. Quantification of phagocytosis index, Aβ_1-42_ oligomer stimulation increased the phagocytosis index compared to vehicle (Veh; Wilcoxon signed rank test, *p* = 0.002). Each dot in the graph represents the mean of three cell culture wells per donor. Data are presented as mean ± SEM. c. Principal component analysis of unstimulated and Aβ_1-42_ oligomer stimulated MDMi samples. d. Volcano plot showing differentially expressed genes when comparing stimulated and unstimulated MDMi cells. e. TPM-normalized read counts of a selection of significantly upregulated genes after Aβ_1-42_ oligomer stimulation. f. Gene ontology and Panther pathway analyses performed on upregulated genes, adj. *p*-value < 0.05 and Log2 FC > 0.5 in MDMi cells after Aβ_1-42_ oligomer stimulation. The number of genes that cluster in the specific terms are shown within the bars.

A PCA plot showed that the Aβ_1-42_ oligomer-stimulated and unstimulated samples (vehicle) do not form separate clusters. Instead, samples from the same donor clustered together (Fig. 3c), indicating a larger difference between donors than induced by the Aβ treatment. Differential gene expression analysis identified 20 genes that were upregulated and two genes that were downregulated (with adj. *p* < 0.05 and a Log2 FC > 0.5) in Aβ_1-42_ oligomer-stimulated MDMi samples (Fig. 3d and Supplementary Table 3). Several subtypes of the *MT1* isoform were upregulated after Aβ_1-42_ oligomer stimulation (Fig. 3e). The top five upregulated genes with the highest log2 FC were all subtypes of metallothionein 1 (*MT1*), a family of cysteine-rich proteins. GO on the 20 differentially expressed genes between vehicle and Aβ_1-42_ oligomer-stimulated MDMi cells identified changes in “metal ion binding”, “transition metal ion binding”, and “zinc ion binding” (Fig. 3f). The Panther pathway analysis suggested changes in “hypoxia and oxidative stress response”, and in the “AD presenilin pathway”. However, these pathways were identified based on the upregulation of only one gene, *viz. TXN* and *CD44*, respectively. The top five upregulated transcripts (*MT1, MT2, SOD2, AKR1B1*, and *C15ORF48*) were validated by qPCR (Supplementary Fig. S1). These results indicated that the transcriptomic changes induced by human Aβ_1-42_ oligomers are predominantly related to metal binding and control of oxidative stress.

### Aβ_1-42_ oligomer response in human primary microglia

Four subtypes of *MT1* (*MT1E, MT1G, MT1L*, and *MT1X*) that are in the top five upregulated genes after Aβ_1-42_ oligomer stimulation in MDMi cells were shown by Bossers et al., 2010 to increase with disease progression and were highest in end-stage AD (Bossers et al., 2010). However, the gene expression profile from Bossers et al., 2010 was not specific to the microglia population (Bossers et al., 2010). Therefore, to determine whether the expression of *MT1, MT2, SOD2, AKR1B1*, and *C15ORF48* is increased in pMG from AD-cases, a qPCR analysis was done on primary microglia isolated from AD-cases microglia and NDC. *MT1* expression was significantly increased in AD-cases (Fig. 4a). The expression of a selection of genes associated with late-onset AD (*CD33, TREM2, TYROBP, ITGAM*, and *ApoE*) was not increased in AD-cases (Fig. S2).

**Figure 4.**
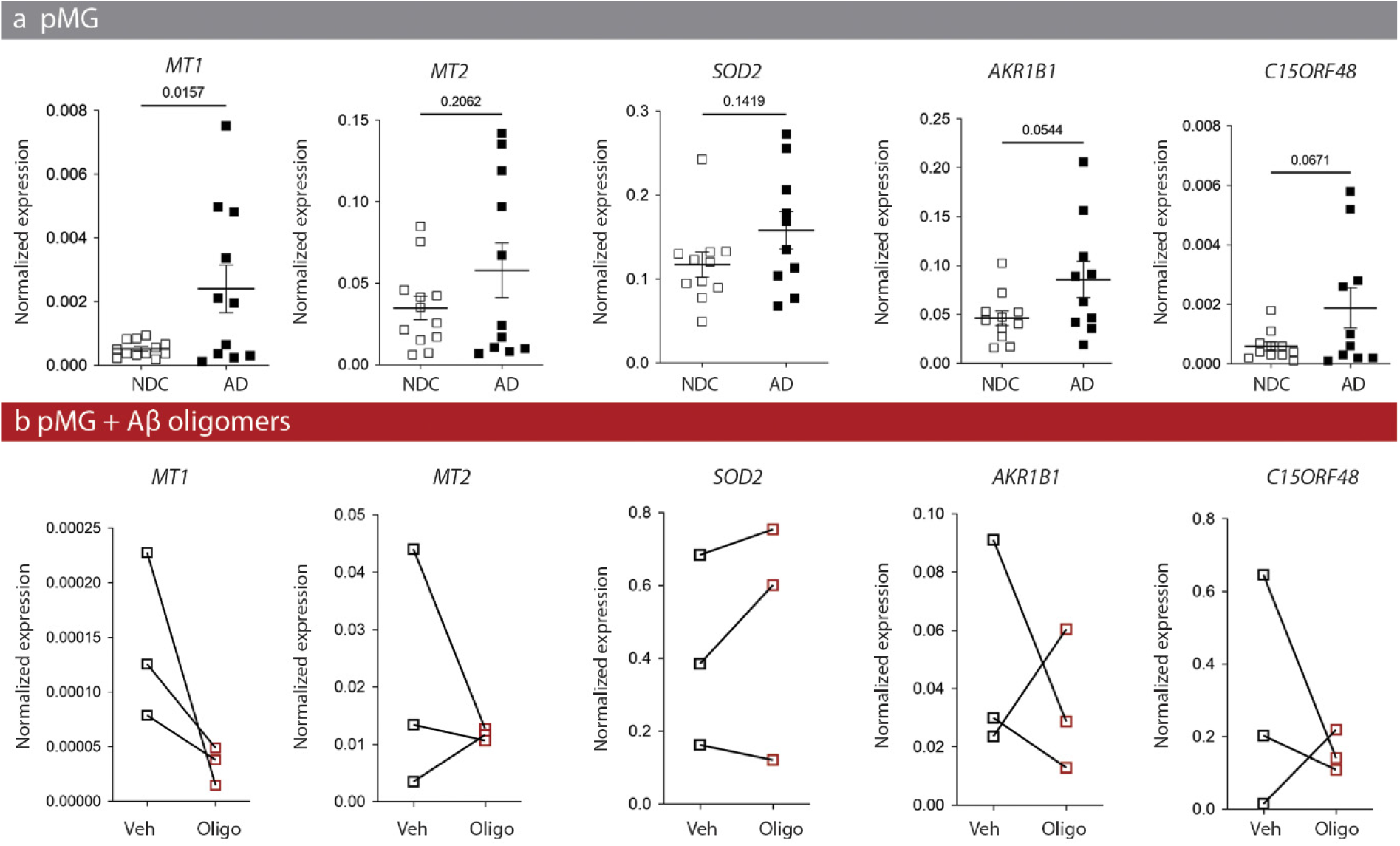
Expression of the **Aβ**_**1-42**_ oligomer profile in isolated microglia. a. Human microglia were isolated from the gyrus temporalis superior (GTS) of NDC-(N = 12) and AD-cases (N = 11). After isolation the mRNA expression of the top five most significantly upregulated genes was determined using qPCR. mRNA expression was normalized to *GAPDH, S18*, and *ACTB*. Data represented as mean ± SEM. Outliers were removed using ROUT test, in *AKR1B1* NDC and AD; *SOD2* NDC and AD; *C15ORF48* NDC and AD. b. mRNA expression of the top five most significantly upregulated genes determined by qPCR following 24 hrs stimulation of microglia (NDC, N = 3) with Aβ_1-42_ oligomers.

To determine whether Aβ_1-42_ oligomer stimulation in pMG recapitulated the gene expression profile found in MDMi cells, a selection of pMG isolated from NDC-cases was cultured for 24-48 hrs and subsequently stimulated with Aβ_1-42_ oligomers for 24 hrs. This stimulation did not induce an upregulation of *MT1, MT2, SOD2, AKR1B1*, and *C15ORF48* (Fig. 4b).

### The Aβ_1-42_ oligomer-induced expression profile in MDMi cells is not detected in isolated microglia of APPswePS1dE9 mice

A previously published dataset showed that *Mt1* (0.8 Log2 FC), *Mt2* (1.4 Log2 FC), and *Sod2* (0.8 Log2 FC) were increased in cortical microglia isolated from 15- to 18-month-old APPswePS1dE9 mice (Orre et al., 2014). We next determined whether these upregulated genes were also increased in cortical microglia from 4-month-old APPswePS1dE9 mice, when increased levels of amyloid and the first plaques are detected (Garcia-Alloza et al., 2006; Van Tijn et al., 2012). qPCR analysis showed no increase in mRNA levels of these genes in 4-month-old APPswePS1dE9 mice (Fig. S3).

### AD pathology-associated microglia profiles

Next, we compared our stimulated and unstimulated MDMi transcriptome datasets to three well-known AD microglia profiles (Keren-Shaul et al., 2017; Mathys et al., 2019; Srinivasan et al., 2019). The studies by Srinivasan *et al*., 2019 and Mathys *et al*., 2019 investigated transcriptome changes in microglia of healthy control donors and AD patients (Mathys et al., 2019; Srinivasan et al., 2019). The first study refers to their novel human Alzheimer’s microglia/myeloid cells profile as the HAM-profile (Srinivasan et al., 2019). The HAM-profile contains 66 differentially expressed genes between AD and control subjects, from which we detected 50 genes in the expression dataset of Aβ_1-42_ oligomer stimulation and 51 genes in the expression dataset of LPS stimulated MDMi cells. However, only *Kcnj5* was differentially expressed in both the HAM signature and our dataset from Aβ_1-42_ oligomer-stimulated MDMi cells (Fig. 5a). In our dataset, this gene was downregulated instead of upregulated (Fig. 5b). The comparison of the HAM signature and our LPS-stimulated MDMi cells identified 20 genes with a p-value < 0.05, of which eight genes were upregulated and one was downregulated in both HAM and LPS-stimulated MDMi cells (Fig. 5b). Overall, the HAM signature is not present in our MDMi cell model after Aβ_1-42_ oligomer stimulation (Fig. 5c). Only a small overlap was detected in the LPS-stimulated MDMi cells (Fig. 5a-b).

**Figure 5.**
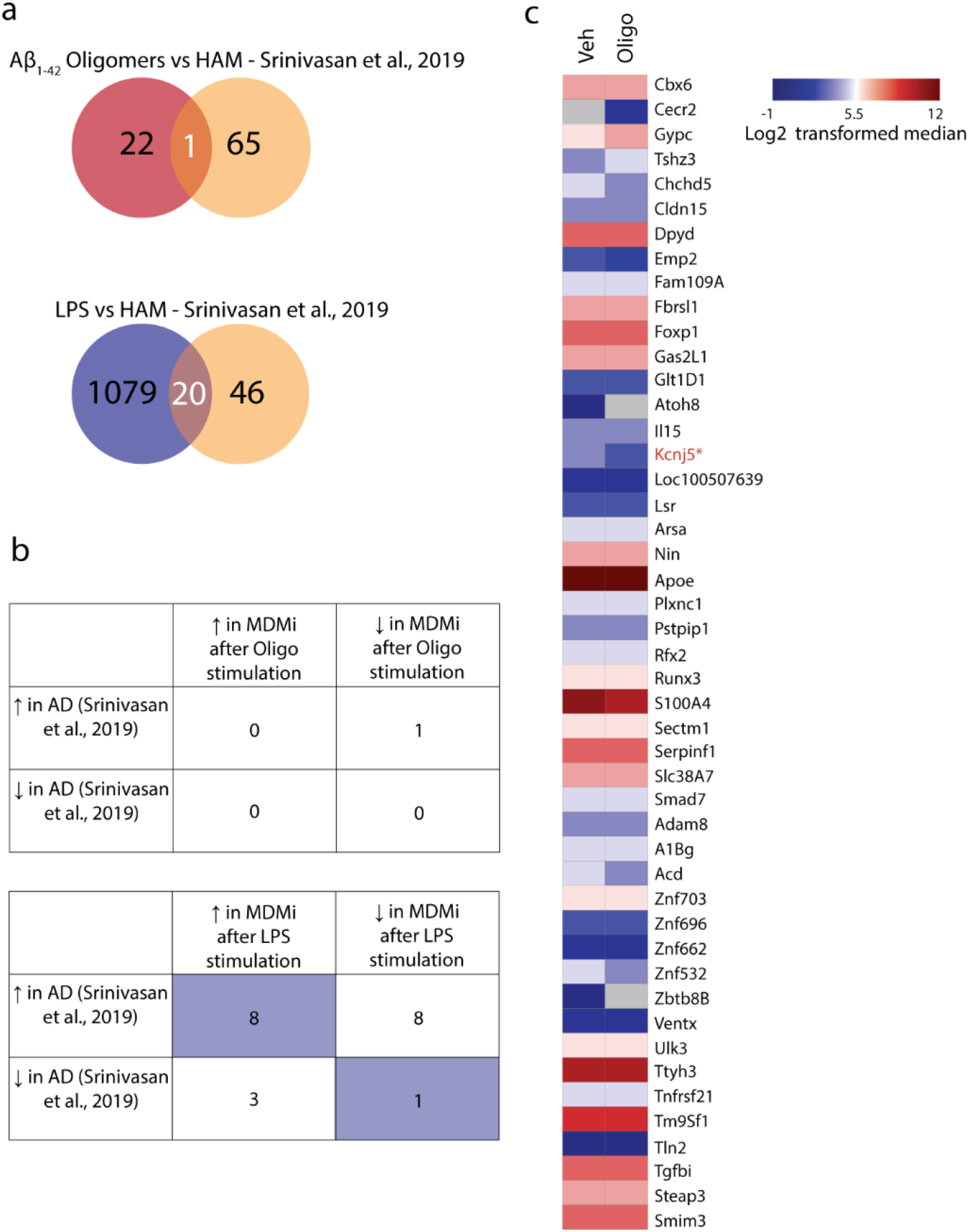
Comparison between the human Alzheimer microglia/myeloid (HAM) profile and Aβ_**1-42**_ oligomer- or LPS-stimulated MDMi cells. a. Venn diagrams showing overlap of the HAM-profile (Srinivasan et al., 2019) in Aβ_1-42_ oligomer- or LPS-stimulated MDMi samples. b. Tables show the number of genes that overlap between the HAM-profile and Aβ_1-42_ oligomer-(top) or LPS-stimulated MDMi cells (bottom). c. Heatmap showing 66 genes from the HAM-profile in vehicle and Aβ_1-42_ oligomer-stimulated MDMi cells. The Log2 transformed median of each group was used for the heatmap.

The AD-pathology associated microglia profile described by Mathys *et al*., 2019 contains 77 differentially expressed genes (Mathys et al., 2019). From this profile, two genes, *Acsl1* and *Fth1*, were also upregulated in Aβ_1-42_ oligomer-stimulated MDMi cells (Fig. S4a-b). In the LPS-stimulated MDMi cells 19 genes from the 77 differentially expressed genes were upregulated. Overall, the AD-pathology associated microglia signature is not present in our MDMi cell model after Aβ_1-42_ oligomer stimulation. Only a partial overlap was detected in the LPS-stimulated MDMi cells (Fig. S4c-d).

The study by Keren-Shaul *et al*., 2017 identified the so-called disease-associated microglia (DAM), in the 5xFAD AD mouse model (Keren-Shaul et al., 2017). The DAM profile is to a certain degree conserved in mice and humans (Mathys et al., 2019). The comparison of the DAM profile and our Aβ_1-42_ oligomer-stimulated MDMi cells identified 17 genes with a p-value < 0.05, from which nine were upregulated and three downregulated in both DAM and Aβ_1-42_ oligomer-stimulated MDMi cells.

## Discussion

In this study, we performed bulk RNA sequencing on unstimulated and stimulated MDMi cells to investigate the effect of LPS and human Aβ_1-42_ oligomers on their transcriptome. We first characterized the MDMi cell model on morphology and gene expression and concluded that they recapitulate some of the key aspects of microglial phenotype and function. In accordance with previous findings, LPS stimulation induced an immune activated phenotype in the MDMi cells (Ormel et al., 2020). Both LPS and Aβ_1-42_ oligomer stimulation increased the phagocytosis index of MDMi cells. Furthermore, the Aβ_1-42_ oligomer stimulation resulted in the upregulation of 20 genes, including several *MT1* subtypes, and the downregulation of two genes. Overall, Aβ_1-42_ oligomers induced less changes in MDMi cells than the strong inflammatory stimulus LPS.

Some of the classical AD markers as identified in GWAS studies, including *CD33, TREM2*, and *ApoE*, were not upregulated by Aβ_1-42_ oligomer stimulation in MDMi cells. Differentially expressed genes from stimulated MDMi samples showed little overlap with AD microglial profiles described in recent transcriptome studies (Keren-Shaul et al., 2017; Mathys et al., 2019; Srinivasan et al., 2019). These findings indicate that stimulation with Aβ_1-42_ oligomers induces a distinct gene expression profile in MDMi cells, different from the gene profile of microglia from end-stage AD pathology, characterized by a high amyloid burden, and, in humans, also increased neurofibrillary tangles. The profile identified in MDMi cells is specific for Aβ_1-42_ oligomer stimulation and not influenced by the effect of other cells in the central nervous system, previous activation of the cells, tau pathology, and/or other amyloid isoforms. Whether an end-stage AD pathology profile could be induced in MDMi cells by co-culturing with other cells, stimulation with tau and/or other amyloid isoforms should be investigated to determine the synergistic effect of these factors.

Several subtypes of a specific family of metalloproteins, the metallothioneins (MTs), were upregulated after stimulation with Aβ_1-42_ oligomers. MTs are low molecular weight cysteine-rich metal-binding proteins. MTs consist of four subfamilies, with MT1 and MT2 being the most predominant isoforms present in most tissues (Vašák and Meloni, 2011). In the central nervous system, MT1 and MT2 are predominantly expressed by astrocytes. However, expression in neurons, endothelial cells, and microglia has also been shown (Pedersen et al., 2009). Although the primary role of MTs remains unclear, increasing evidence indicates that these proteins have multiple functions, including maintenance of zinc and copper homeostasis, anti-inflammatory properties, regulating the biosynthesis and activity of zinc-binding proteins, and protection against reactive oxygen species (Hozumi et al., 2004; Manso et al., 2012; Waller et al., 2018). Their expression is increased in response to a variety of stimuli, including oxidative stress, neuroinflammation, and toxic levels of metal ions (West et al., 2008). In mice, it was found that *MT1* and *MT2* were upregulated in the brain five hrs after LPS injection (Searle et al., 1984). Also in MDMi cells stimulated with LPS, an upregulation of several *MT1* subtypes and *MT2A* was found. In several neurodegenerative disorders, including AD, MT1 and MT2 are upregulated (Adlard et al., 1998; Duguid et al., 1989; Zambenedetti et al., 1998) and associated with Aβ plaques in several AD animal models (Carrasco et al., 2006; Hidalgo et al., 2006). The upregulation of MTs could have a neuroprotective function against the oxidative stress and neuroinflammation involved in AD pathogenesis (Adlard et al., 1998; Nunomura et al., 2006). Also, some other genes encoding for proteins known for their anti-oxidant response were upregulated in the Aβ_1-42_ oligomer stimulated MDMi, such as SOD2 (Massaad et al., 2009) and NQO1 (Raina et al., 1999; SantaCruz et al., 2004).

Bossers *et al*., 2010 showed that the expression of several *MT1* isoforms is upregulated during the progression of AD. However, this gene expression profile was obtained from the grey matter of the frontal cortex and was not specific to the microglia population (Bossers et al., 2010). Interestingly *MT1* was also significantly upregulated in isolated microglia from AD-compared to NDC-cases. Based on the mRNA expression of both *MT2* and *C15ORF48*, two groups of donors seem to be present within the AD-cases. However, no significant association was detected with the known confounding variables age, sex, *post-mortem* delay, pH of the cerebral spinal fluid or with Braak or amyloid scores (Supplementary Table 5 and 6). The expression of a selection of genes associated with late-onset AD (*CD33, TREM2, TYROBP, ITGAM*, and *ApoE*) were not upregulated in the isolated microglia of AD-cases. Although several recent studies found differences between the gene expression profile of microglia from AD- and NDC-cases (Grubman et al., 2019; Mathys et al., 2019; Srinivasan et al., 2019), this was not found in all studies (Alsema et al., 2020).

Culturing primary microglia from a selection of NDC-cases for 24-48 hrs and subsequent 24 hrs stimulation with Aβ_1-42_ oligomers did not induce the upregulation of the top upregulated genes detected in MDMi cells. This is in contrast with the study by Walker *et al*., 2006 where stimulation for 24 hrs with oligomeric amyloid resulted in the activation of an inflammatory response and the upregulation of *MT1, MT2* and *SOD2* in cultured microglia from the superior frontal cortex (Walker et al., 2006). The genes were categorized as anti-inflammatory and anti-oxidant proteins. Besides the upregulation of these three genes, *KYNU*, also present in the Aβ_1-42_ oligomer profile, was increased as well (Walker et al., 2006). Differences in the concentration of Aβ_1-42_ oligomers (*i*.*e*. 500 nM or 2 μM) used, isolation procedures or the duration of the microglia in culture prior to the stimulation (*i*.*e*. 24-48 hrs or 12-24 days) could explain the different response to Aβ_1-42_ oligomer stimulation. We waited 48 hrs before Aβ_1-42_ oligomer stimulation, which might have been too short to bring the microglia to a nonactivated state after isolation. However, as microglial markers are downregulated in a culture environment (Gosselin et al., 2017), culturing the microglia for a longer period also introduces other limitations.

In summary, in this study we investigated the transcriptomic changes following stimulation of MDMi cells with inflammatory stimulus and Aβ_1-42_ oligomers. LPS stimulation induced an immune activated phenotype, and stimulation with stable Aβ_1-42_ oligomers induced a specific more anti-inflammatory transcriptome profile in MDMi cells. Several anti-oxidant genes were upregulated by Aβ_1-42_ oligomer stimulation, most notably metallothionein subtypes, which are known to be involved in metal ion regulation, protection against reactive oxygen species and have anti-inflammatory properties. In conclusion, the role of metallothioneins in AD pathogenesis and their induced upregulation after acute exposure to Aβ_1-42_ oligomers should be investigated further to determine whether the acute activation by oligomeric amyloid could trigger a protective response.

## Acknowledgements

This work was supported by a TAS-ZonMw grant (40-41400-98-16020), ZonMw grant (733050505) and a Memorabel-ZonMw grant (733050816) to E.M.H. We are grateful to the Netherlands Brain Bank (www.brainbank.nl) who provided us with the *post-mortem* human brain tissue. We are also grateful to Guus Scheefhals from Crossbeta for providing the Aβ_1-42_ oligomers used in this study. We thank Y. He, M. P. Boks, and M. Litjens for the isolation of monocytes, C. San Martin Paniello for assistance with setting up the MDMi cell model, and R.D. van Dijk and M.A.M. Sneeboer for their help with setting up the microglial isolations and phagocytosis assays using FACS.

## Supplementary data

**Supplementary Table 1.** Overview of donors used for MDMi differentiation and transcriptome analysis. **\*** Samples that had more than 10M reads were discarded. F = female, M = male. Average age: 51 ± 12 (standard deviation).

| Donor | Condition | Library | Primer used | Number of reads | Sex | Age (years) |
| --- | --- | --- | --- | --- | --- | --- |
| M | PBS | UMC-TS-B001 | I1 | 4670509 | F | - |
| M | LPS | UMC-TS-B001 | I3 | 5997461 |  |  |
| M | Vehicle | UMC-TS-B001 | I5 | 4550292 |  |  |
| M | Oligomers | UMC-TS-B001 | I7 | 4022624 |  |  |
| A | PBS | UMC-TS-B002 | I9 | 4162021 | F | 50 |
| A | LPS | UMC-TS-B002 | I11 | 4326116 |  |  |
| A | Vehicle | UMC-TS-B002 | I13 | 2334147 |  |  |
| A | Oligomers | UMC-TS-B002 | I15 | 3685513 |  |  |
| B | PBS | UMC-TS-B003 | I17 | 4586612 | M | 62 |
| B | LPS | UMC-TS-B003 | I19 | 7390755 |  |  |
| B | Vehicle | UMC-TS-B003 | I21 | 4315275 |  |  |
| B | Oligomers | UMC-TS-B003 | I23 | 4639184 |  |  |
| C | PBS | UMC-TS-B004 | J1 | 3666645 | F | 27 |
| C | LPS | UMC-TS-B004 | J3 | 8689959 |  |  |
| C | Vehicle | UMC-TS-B004 | J5 | 3291413 |  |  |
| C | Oligomers | UMC-TS-B004 | J7 | 10534576* |  |  |
| AG | PBS | UMC-TS-B005 | J9 | 7851479 | M | 46 |
| AG | LPS | UMC-TS-B005 | J11 | 7275576 |  |  |
| AG | Vehicle | UMC-TS-B005 | J13 | 9068876 |  |  |
| AG | Oligomers | UMC-TS-B005 | J15 | 7285752 |  |  |
| AK | PBS | UMC-TS-B006 | J17 | 13380240* | M | 46 |
| AK | LPS | UMC-TS-B006 | J19 | 3006578 |  |  |
| AK | Vehicle | UMC-TS-B006 | J21 | 5635157 |  |  |
| AK | Oligomers | UMC-TS-B006 | J23 | 3073040 |  |  |
| AA | PBS | UMC-TS-B007 | K1 | 2983729 | F | 54 |
| AA | LPS | UMC-TS-B007 | K3 | 6084654 |  |  |
| AA | Vehicle | UMC-TS-B007 | K5 | 5409584 |  |  |
| AA | Oligomers | UMC-TS-B007 | K7 | 6195652 |  |  |
| AJ | PBS | UMC-TS-B008 | K9 | 7165526 | F | 60 |
| AJ | LPS | UMC-TS-B008 | K11 | 6374208 |  |  |
| AJ | Vehicle | UMC-TS-B008 | K13 | 1598263 |  |  |
| AJ | Oligomers | UMC-TS-B008 | K15 | 5801860 |  |  |
| AB | PBS | UMC-TS-B009 | K17 | 4561945 | F | - |
| AB | LPS | UMC-TS-B009 | K19 | 4113675 |  |  |
| AB | Vehicle | UMC-TS-B009 | K21 | 2728923 |  |  |
| AB | Oligomers | UMC-TS-B009 | K23 | 3799931 |  |  |
| Z | PBS | UMC-TS-B010 | L1 | 2298266 | M | 64 |
| Z | LPS | UMC-TS-B010 | L3 | 2227539 |  |  |
| Z | Vehicle | UMC-TS-B010 | L5 | 3195964 |  |  |
| Z | Oligomers | UMC-TS-B010 | L7 | 1606654 |  |  |

**Supplementary Table 2.**
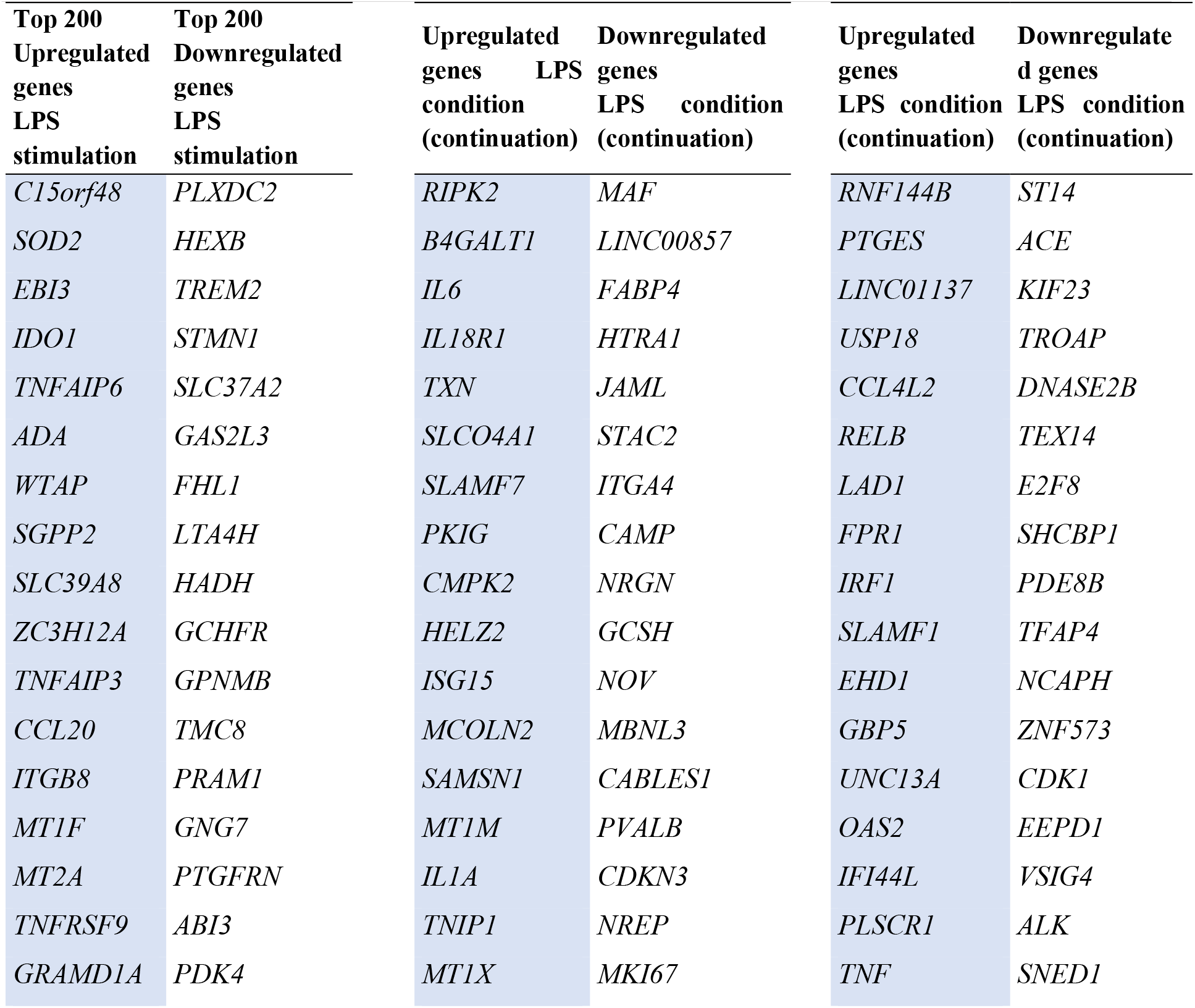
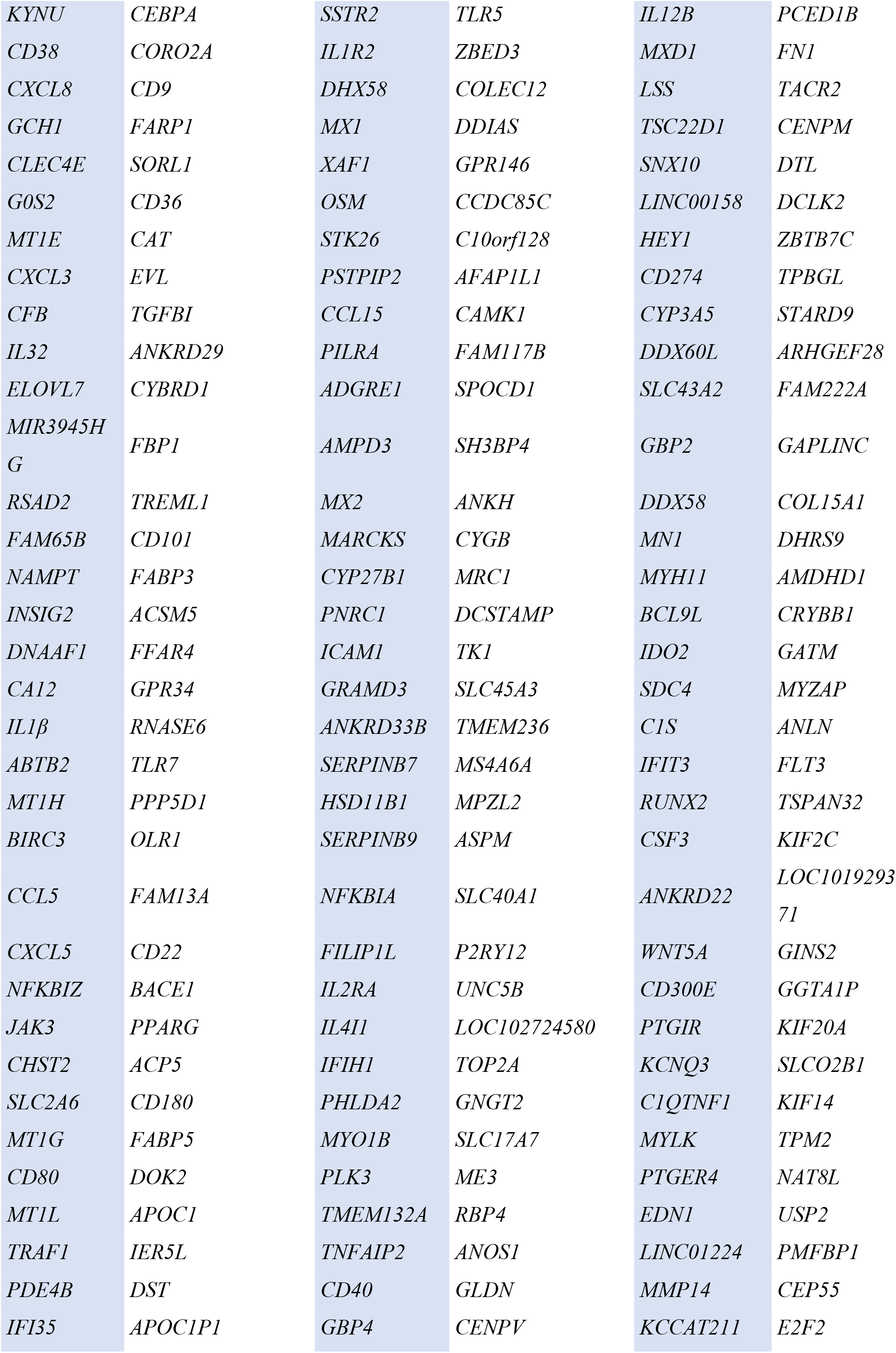
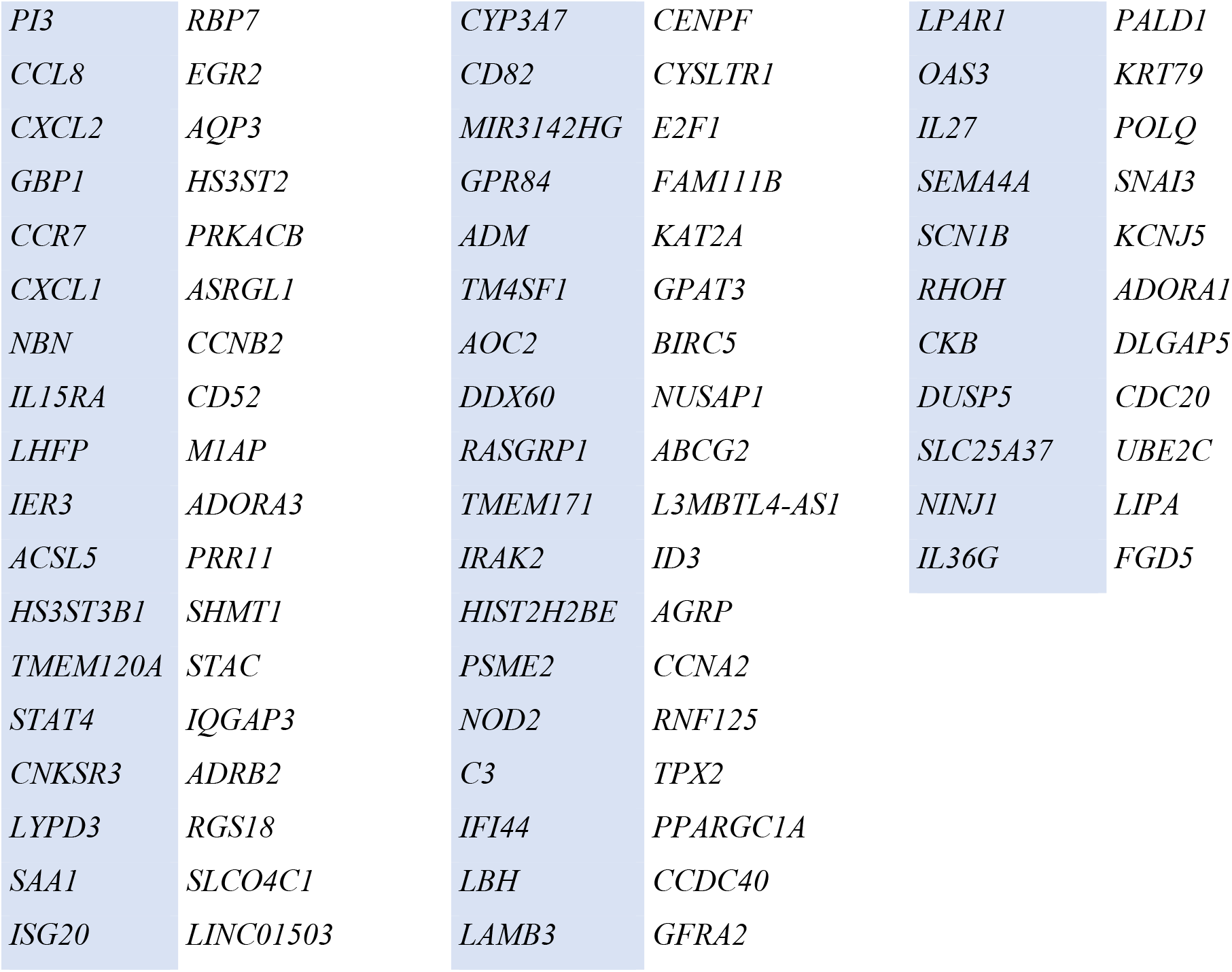
Genes used for the gene ontology (GO) analysis (GO Molecular Function 2018) and Panther pathway analysis (Panther 2016).

**Supplementary Table 3.**
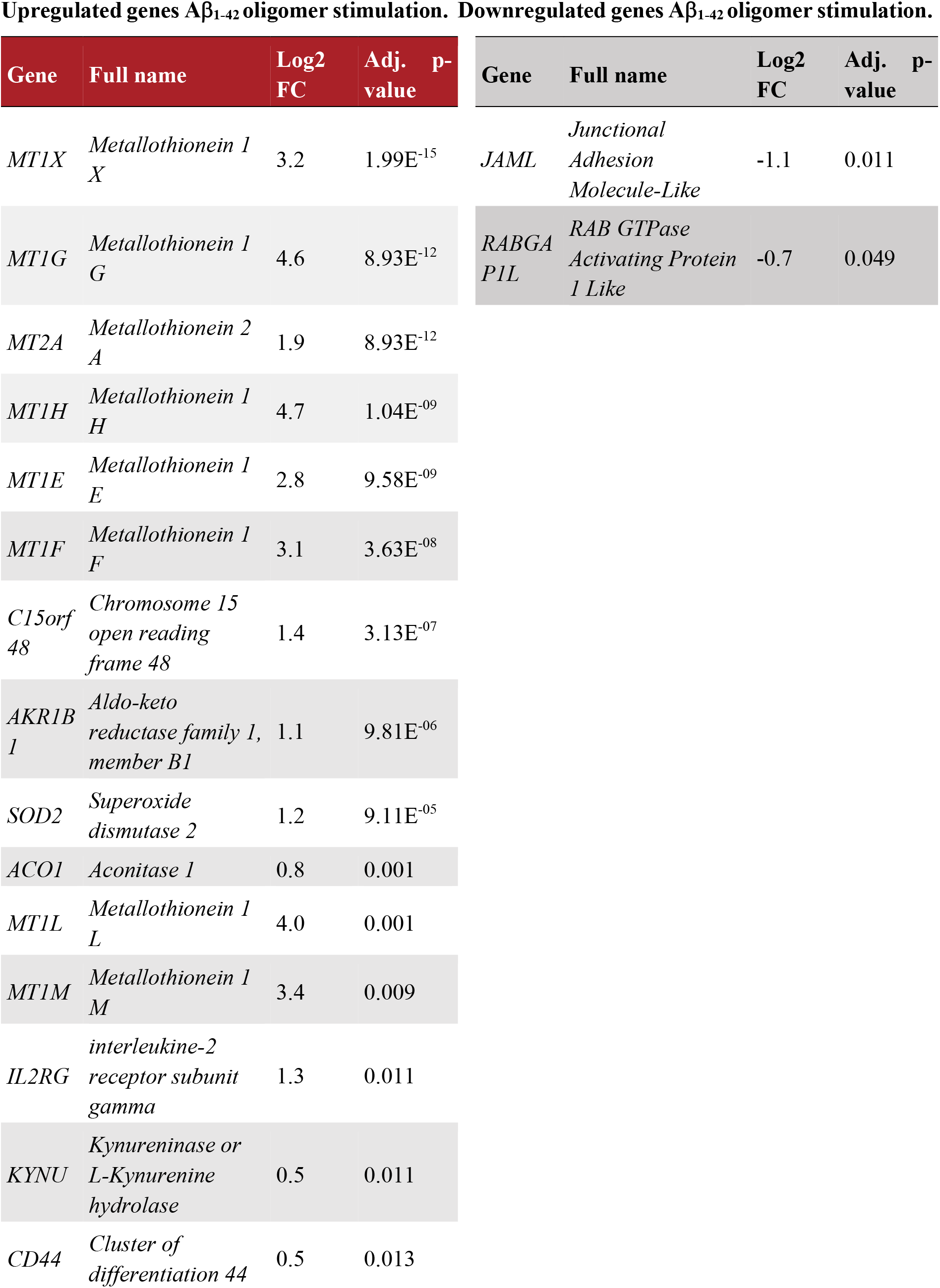
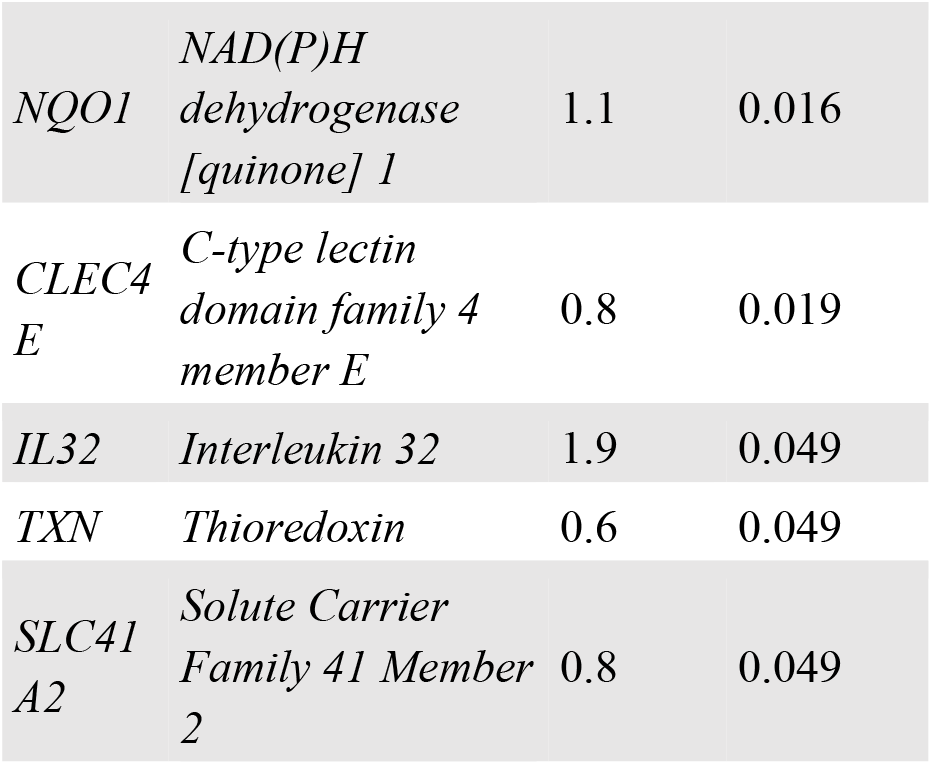
Genes used for the gene ontology (GO) analysis (GO Molecular Function 2018) and Panther pathway analysis (Panther 2016)

**Supplementary Table 4.**
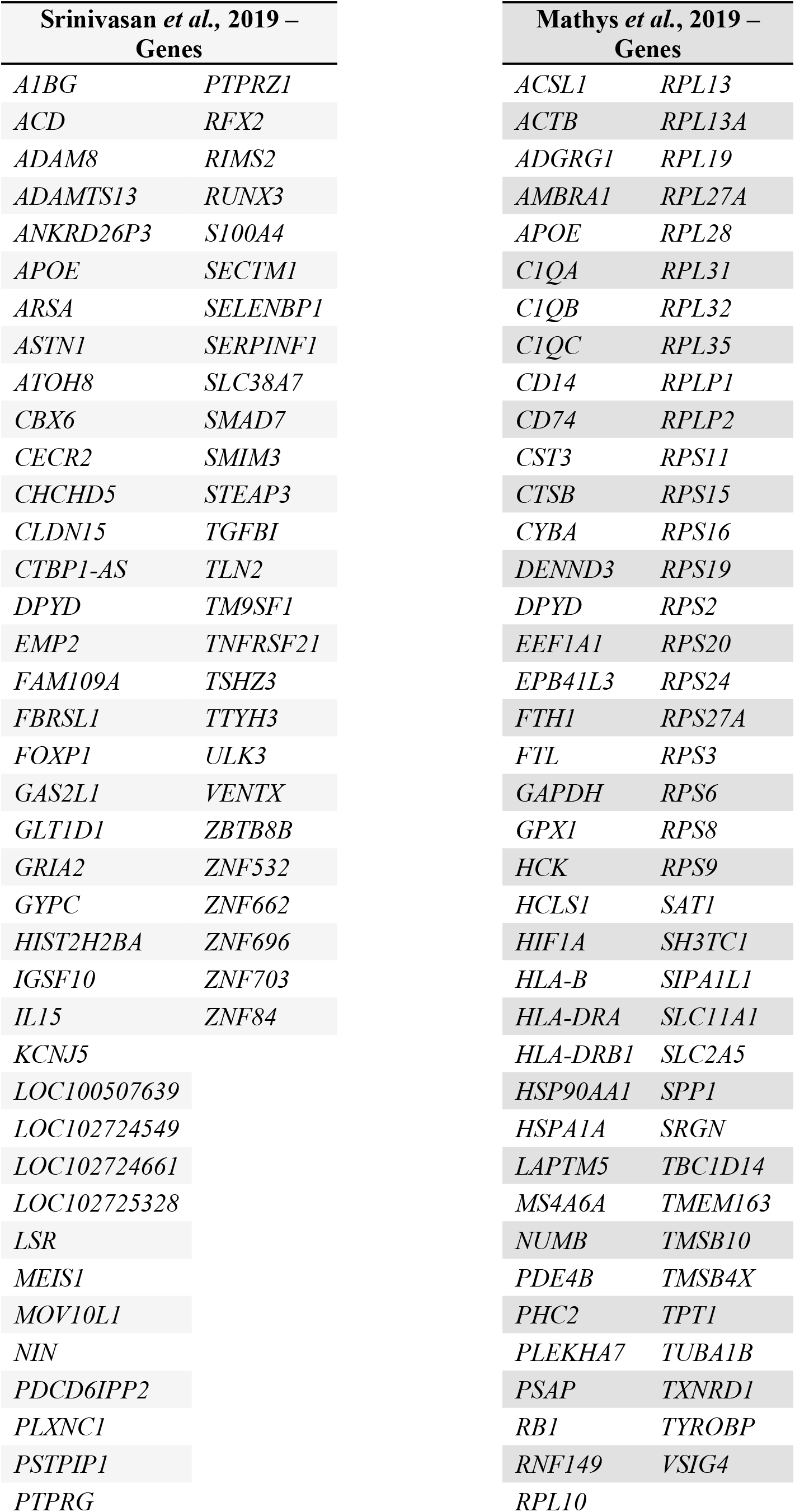
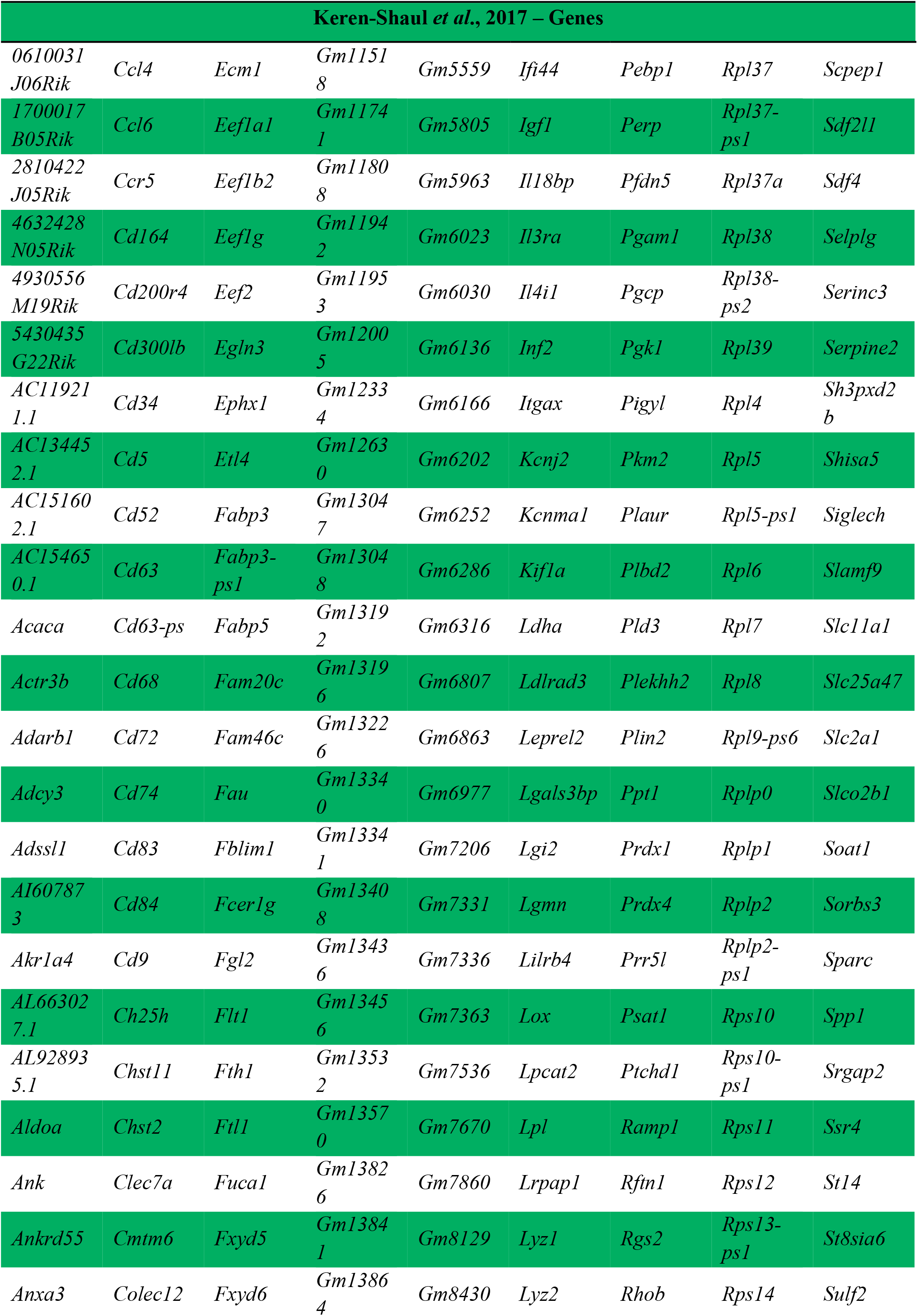
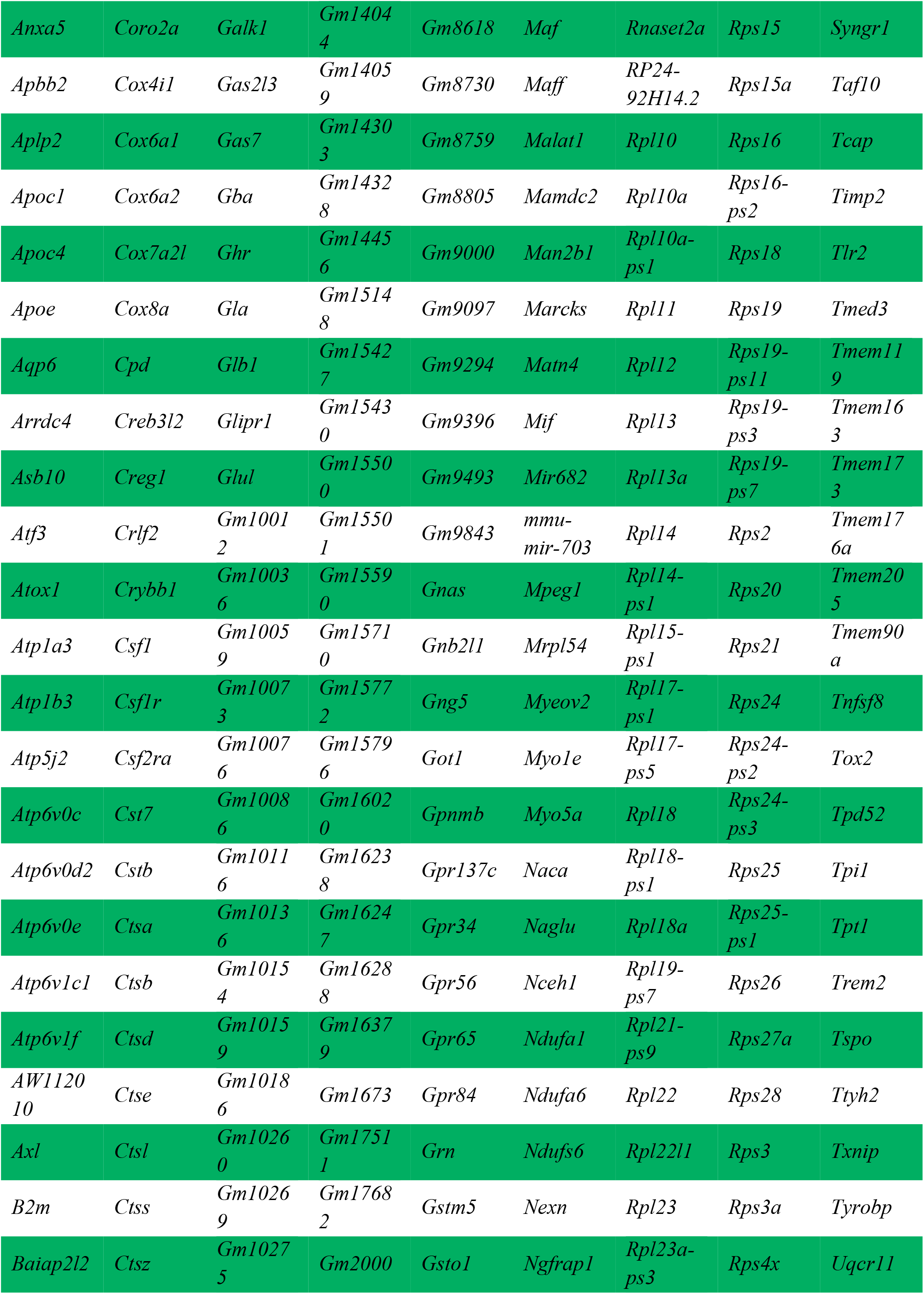
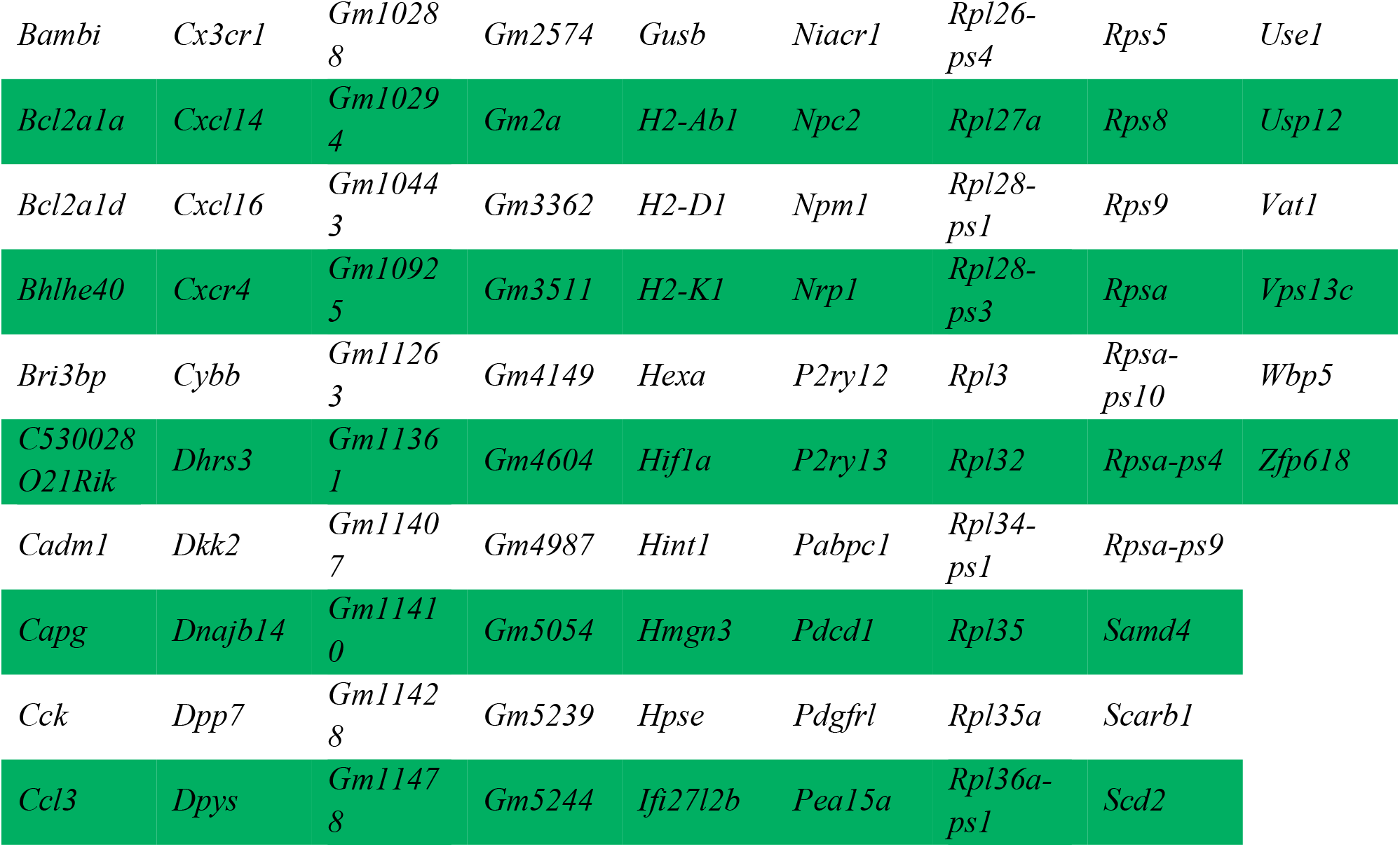
Comparison with previously published datasets.

**Supplementary Table 5.** Clinicopathological information of donors included in this study. Sex, age, *post-mortem* delay (PMD), pH cerebrospinal fluid (CSF), Braak score, amyloid score and brain weight information of cases used in the study. AD = Alzheimer’s disease, F = females, M = males, NDC = non-demented controls.

| NBB Number | Used for figures | Sex | Age (years) | PMD (h:min) | pH CSF | Braak | ABC score | Brain weight (g) |
| --- | --- | --- | --- | --- | --- | --- | --- | --- |
| NDC: |  |  |  |  |  |  |  |  |
| 2018-018 | 4a, S2a | F | 82 | 05:30 | 6.48 | 2 | A0B1C0 | 1165 |
| 2018-030 | 4a, S2a | F | 102 | 05:10 | 6.52 | 4 | A1B2C0 | 1093 |
| 2018-105 | 4, S2 | M | 86 | 06:45 | 6.39 | 3 | A0B2C0 | 1340 |
| 2018-123 | 4a, S2a | F | 82 | 06:20 | - | 3 | A1B2C0 | 1275 |
| 2018-128 | 4a, S2a | M | 94 | 05:30 | 6.57 | - | - | 1229 |
| 2019-029 | 4a, S2a | M | 87 | 06:21 | 6.47 | 3 | A0B2C0 | 1275 |
| 2019-036 | 4, S2 | F | 85 | 06:50 | 6.84 | 3 | A1B2C0 | 1190 |
| 2019-042 | 4a, S2a | M | 82 | 06:30 | 6.53 | 2 | A0B1C0 | 1270 |
| 2019-050 | 4, S2 | M | 77 | 08:45 | 6.39 | - | - | 1319 |
| 2019-061 | 4a, S2a | M | 82 | 05:55 | 6.05 | 2 | A0B1C0 | 1580 |
| 2019-077 | 4a, S2a | F | 91 | 09:30 | 6.36 | 3 | - | 1015 |
| 2019-079 | 4a, S2a | F | 83 | 07:50 | 6.59 | 3 | A1B2C0 | 1345 |
| AD: |  |  |  |  |  |  |  |  |
| 2018-034 | 4a, S2a | M | 78 | 06:35 | 6.61 | 6 | A3B3C3 | 1088 |
| 2018-048 | 4a, S2a | M | 76 | 04:40 | 6.40 | 6 | A3B3C3 | 1300 |
| 2019-073 | 4a, S2a | M | 96 | 04:50 | 7.33 | 3 | A3B2C2 | 1209 |
| 2018-077 | 4a, S2a | M | 86 | 03:45 | 6.54 | 4 | A3B2C2 | 1406 |
| 2018-080 | 4a, S2a | F | 94 | 04:00 | 6.36 | - | - | 1116 |
| 2018-082 | 4a, S2a | M | 68 | 04:30 | 6.43 | 6 | A3B3C3 | 1163 |
| 2018-090 | 4a, S2a | M | 97 | 05:10 | 6.65 | 5 | A3B3C3 | 1120 |
| 2019-010 | 4a, S2a | F | 76 | 03:45 | 6.55 | 5 | A3B3C3 | 1155 |
| 2019-030 | 4a, S2a | M | 77 | 04:26 | 6.63 | 4 | A2B2C2 | 1514 |
| 2019-032 | 4a, S2a | M | 92 | 05:50 | 6.35 | 4 | A2B2C2 | 1270 |
| 2019-080 | 4a, S2a | F | 80 | 05:25 | 6.79 | 5 | A3B3C3 | 1045 |

**Supplementary Table 6.**
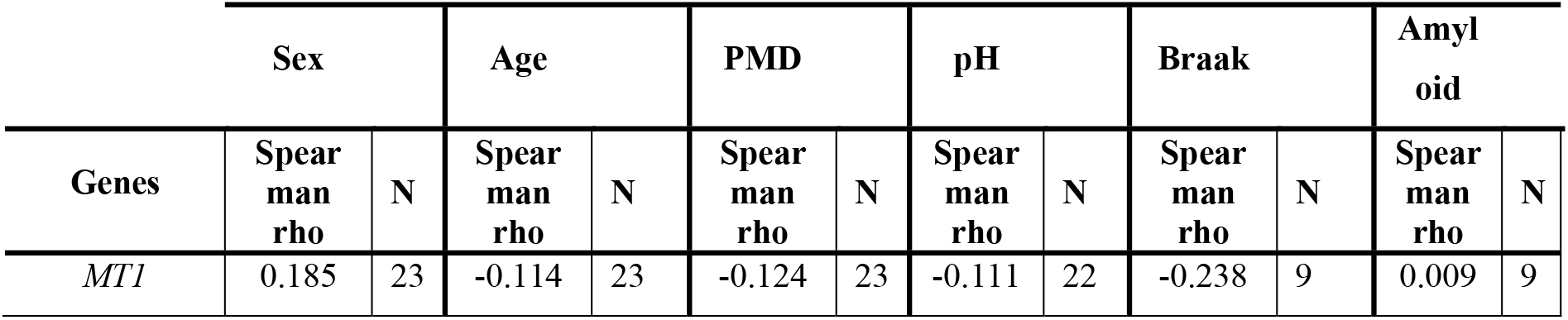
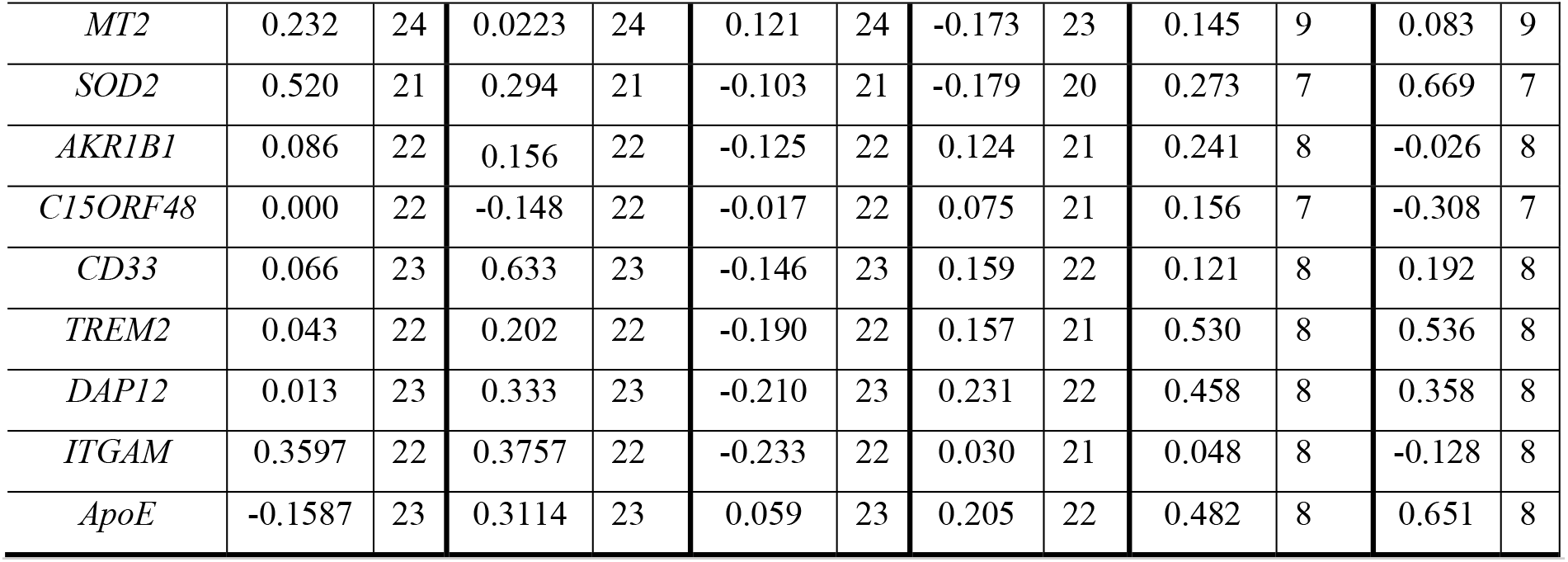
Spearman rho correlations of confounding variables and clinicopathological information. Spearman rho correlations, Spearman rho and number of donors included (N), between the confounding variables and clinicopathological information (sex, age, *post-mortem* delay (PMD), pH of cerebrospinal fluid, Braak, and Amyloid score) and the mRNA expression of isolated microglia of the GTS. No significant values were detected after Bonferroni correction for multiple testing.

**Supplementary Table 7.**
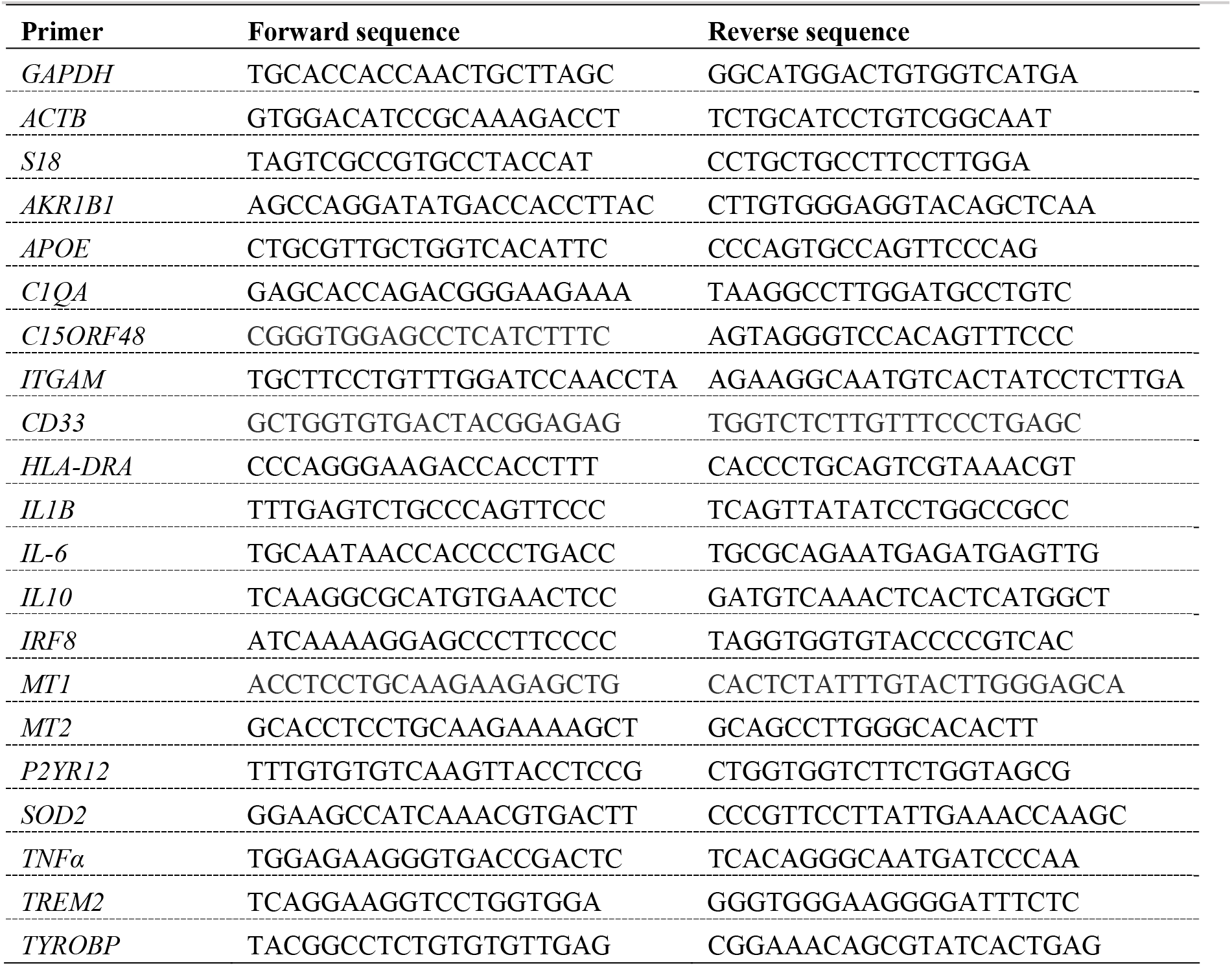
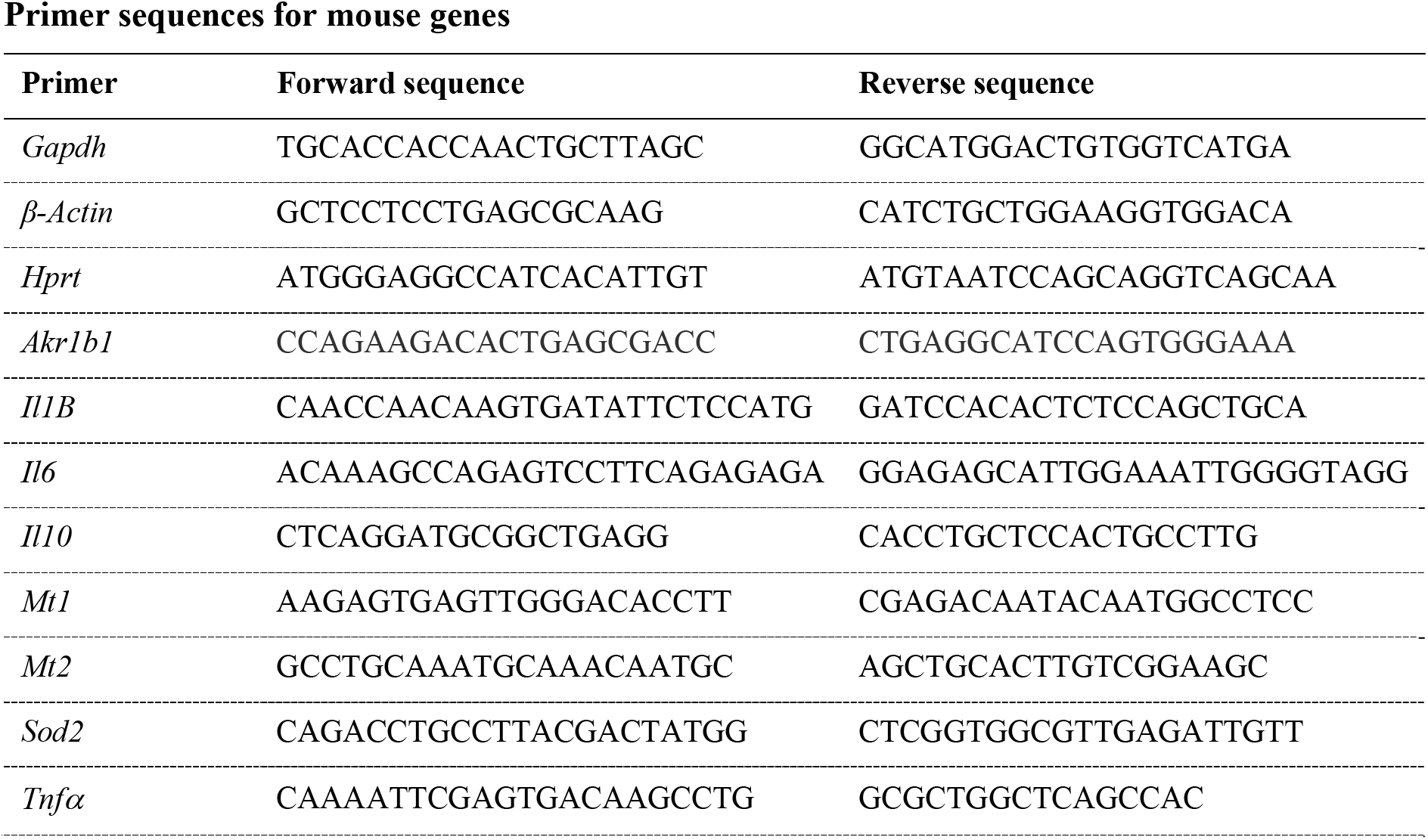
primer sequences.

**Supplementary figure 1.**
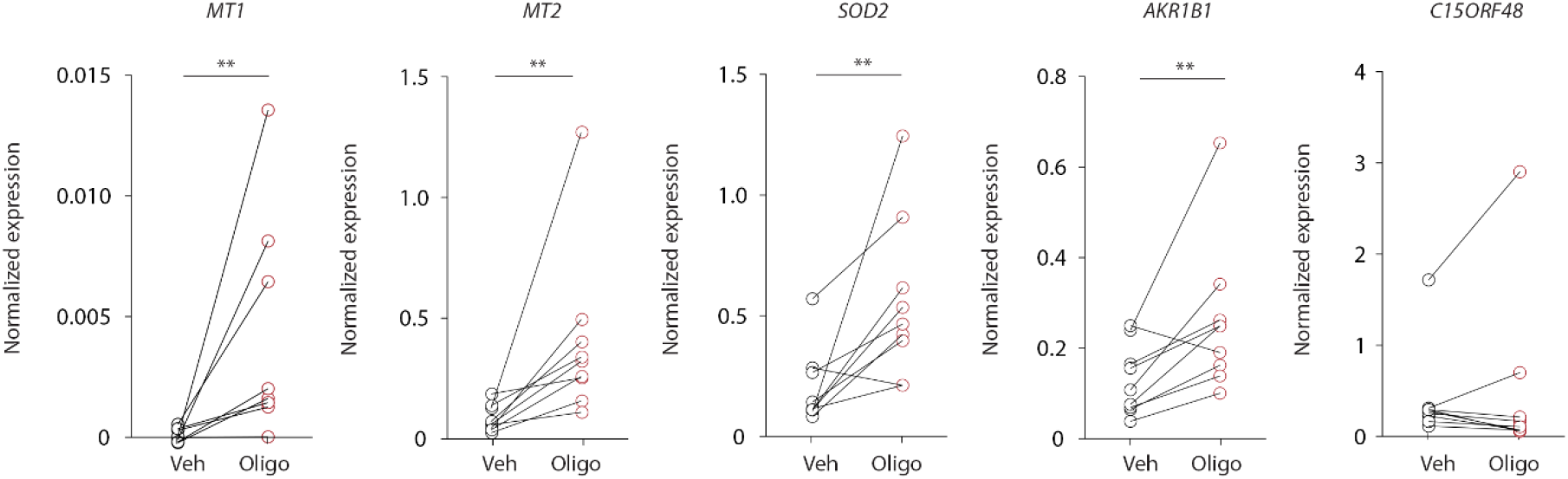
Gene expression of the top five upregulated genes in MDMi cells after Aβ_**1-42**_ oligomer stimulation. mRNA expression of *MT1, MT2, SOD2, AKR1B1*, and *C15ORF48* in unstimulated (vehicle: Veh) and Aβ_1-42_ oligomer-stimulated (Oligo) MDMi cells was determined using qPCR. mRNA expression is normalized to *ACTB, GAPDH*, and *18S*. Wilcoxon signed rank test, *MT1 p* = 0.008, *MT2 p* = 0.004, *SOD2 p* = 0.008, *AKR1B1 p* = 0.008, and *C15ORF48 p* = 0.57.

**Supplementary figure 2.**
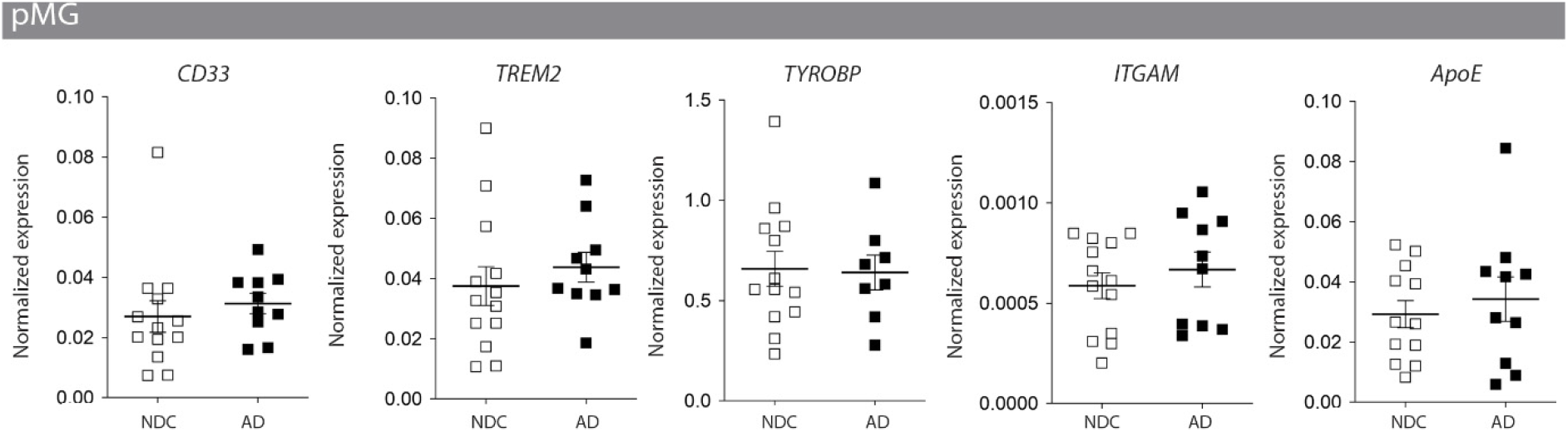
mRNA expression of AD-associated profile and inflammatory markers in isolated microglia from the gyrus temporalis superior (GTS). mRNA expression of a selection of genes associated with late-onset AD in primary microglia isolated from the GTS of NDC-(N = 12) and AD-cases (N = 10) was determined using qPCR. 2 outliers were removed from *DAP12* AD. mRNA expression was normalized to *ACTB, GAPDH*, and *S18*.

**Supplementary figure 3.**
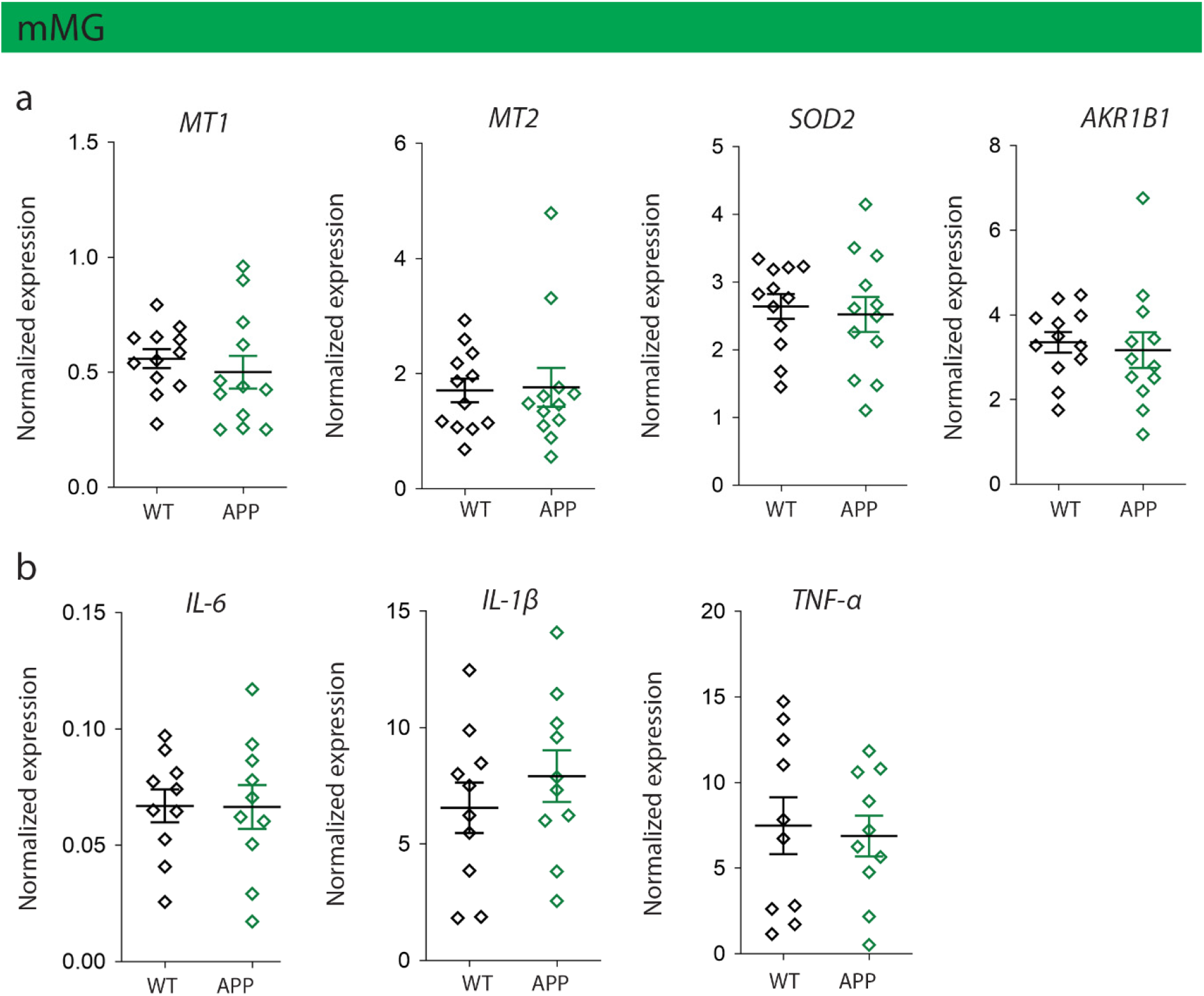
mRNA expression of isolated cortical microglia of APPswePS1dE9 and WT mice. mRNA expression of isolated microglia of the cortex of 4-month-old wild-type littermates (N = 10-12) and APPswePS1dE9 (N = 10-12) was determined by qPCR. a. mRNA expression of four genes of the Aβ_1-42_ oligomer profile, Mann Whitney test, *Mt1 p* = 0.3, *Mt2 p* = 0.8, *Sod2 p* = 0.7, and *Akr1b1 p* = 0.4. b. mRNA expression of a selection of genes associated with an inflammatory response, Mann Whitney test, *Il6 p* = 0.9, *Il1β p* = 0.5, and *Tnfα p* = 0.7. mRNA expression was normalized to reference genes *β-actin, GAPDH*, and *HPRT*. Data are represented as mean ± SEM.

**Supplementary figure 4.**
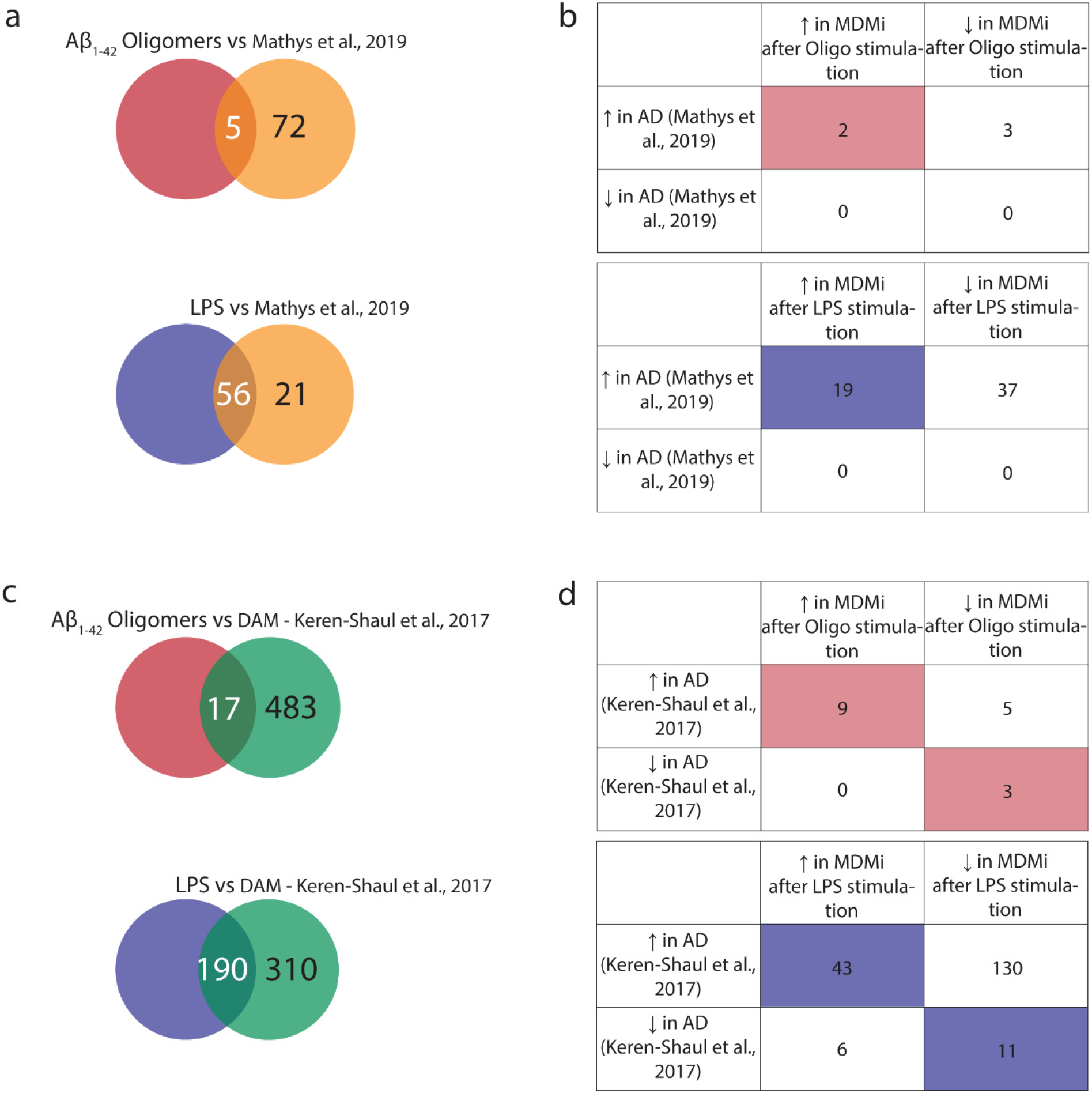
Comparison between the human Alzheimer’s microglia profile obtained by Mathys *et al*., 2019, the DAM-profile, and Aβ_1-42_ oligomer or LPS stimulated MDMi cells. a. Venn diagrams showing overlap of the AD microglia profile (Mathys et al., 2019) in Aβ_1-42_ oligomer and LPS stimulated MDMi samples. b. Tables show the number genes that overlap between the AD microglia profile and Aβ_1-42_ oligomer (top) or LPS stimulated MDMi cells (bottom). c. Venn diagrams showing overlap of the disease-associated microglia (DAM) profile (Keren-Shaul et al., 2017) in Aβ_1-42_ oligomer and LPS stimulated MDMi samples. d. Tables show the number genes that overlap between the DAM profile and Aβ_1-42_ oligomer (top) or LPS stimulated MDMi cells (bottom).

## References

Adlard, P.A., West, A.K., Vickers, J.C., 1998. Increased density of metallothionein I/II-immunopositive cortical glial cells in the early stages of Alzheimer’s disease. Neurobiol. Dis. 5, 349–356. https://doi.org/10.1006/nbdi.1998.0203

Alsema, A.M., Jiang, Q., Kracht, L., Gerrits, E., Dubbelaar, M.L., Miedema, A., Brouwer, N., Woodbury, M., Wachter, A., Xi, H.S., Möller, T., Biber, K.P., Kooistra, S.M., Boddeke, E.W.G.., Eggen, B.J.L., 2020. Profiling microglia from AD donors and non-demented elderly in acute human post-mortem cortical tissue. bioRxiv 2020.03.18.995332. https://doi.org/10.1101/2020.03.18.995332

Bamberger, M.E., Harris, M.E., McDonald, D.R., Husemann, J., Landreth, G.E., 2003. A cell surface receptor complex for fibrillar β-amyloid mediates microglial activation. J. Neurosci. 23, 2665–2674. https://doi.org/10.1523/jneurosci.23-07-02665.2003

Bossers, K., Wirz, K.T.S., Meerhoff, G.F., Essing, A.H.W., Van Dongen, J.W., Houba, P., Kruse, C.G., Verhaagen, J., Swaab, D.F., 2010. Concerted changes in transcripts in the prefrontal cortex precede neuropathology in Alzheimer’s disease. Brain 133, 3699–3723. https://doi.org/10.1093/brain/awq258

Butovsky, O., Weiner, H.L., 2018. Microglial signatures and their role in health and disease. Nat. Rev. Neurosci. 19, 622–635. https://doi.org/10.1038/s41583-018-0057-5

Carrasco, J., Adlard, P., Cotman, C., Quintana, A., Penkowa, M., Xu, F., Van Nostrand, W.E., Hidalgo, J., 2006. Metallothionein-I and-III expression in animal models of Alzheimer disease. Neuroscience 143, 911–922. https://doi.org/10.1016/j.neuroscience.2006.08.054

Chen, E.Y., Tan, C.M., Kou, Y., Duan, Q., Wang, Z., Meirelles, G. V., Clark, N.R., Ma’ayan, A., 2013. Enrichr: Interactive and collaborative HTML5 gene list enrichment analysis tool. BMC Bioinformatics 14. https://doi.org/10.1186/1471-2105-14-128

Doens, D., Fernández, P.L., 2014. Microglia receptors and their implications in the response to amyloid β for Alzheimer’s disease pathogenesis. J. Neuroinflammation 11, 1–14. https://doi.org/10.1186/1742-2094-11-48

Duguid, J.R., Bohmont, C.W., Liu, N., Tourtellotte, W.W., 1989. Changes in brain gene expression shared by scrapie and Alzheimer disease. Proc. Natl. Acad. Sci. U. S. A. 86, 7260–7264. https://doi.org/10.1073/pnas.86.18.7260

Esparza, T.J., Zhao, H., Cirrito, J.R., Cairns, N.J., Bateman, R.J., Holtzman, D.M., Brody, D.L., 2013. Amyloid-beta oligomerization in Alzheimer dementia versus high-pathology controls. Ann. Neurol. 73, 104–119. https://doi.org/10.1002/ana.23748

Garcia-Alloza, M., Robbins, E.M., Zhang-Nunes, S.X., Purcell, S.M., Betensky, R.A., Raju, S., Prada, C., Greenberg, S.M., Bacskai, B.J., Frosch, M.P., 2006. Characterization of amyloid deposition in the APPswe/PS1dE9 mouse model of Alzheimer disease. Neurobiol. Dis. 24, 516–524. https://doi.org/10.1016/j.nbd.2006.08.017

Ginhoux, F., Greter, M., Leboeuf, M., Nandi, S., See, P., Gokhan, S., Mehler, M.F., Conway, S.J., Ng, L.G., Stanley, E.R., Samokhvalov, I.M., Merad, M., 2010. Fate mapping analysis reveals that the adult microglia derive from primitive macrophages. Science (80-.). 330, 841–845. https://doi.org/10.1126/science.1194637

Ginhoux, F., Lim, S., Hoeffel, G., Low, D., Huber, T., 2013. Origin and differentiation of microglia. Front. Cell. Neurosci. 7, 1–14. https://doi.org/10.3389/fncel.2013.00045

Gosselin, D., Skola, D., Coufal, N.G., Holtman, I.R., Schlachetzki, J.C.M., Sajti, E., Jaeger, B.N., O’Connor, C., Fitzpatrick, C., Pasillas, M.P., Pena, M., Adair, A., Gonda, D.D., Levy, M.L., Ransohoff, R.M., Gage, F.H., Glass, C.K., 2017. An environment-dependent transcriptional network specifies human microglia identity. Science (80-.). 356, 1248–1259. https://doi.org/10.1126/science.aal3222

Grubman, A., Chew, G., Ouyang, J.F., Sun, G., Choo, X.Y., McLean, C., Simmons, R.K., Buckberry, S., Vargas-Landin, D.B., Poppe, D., Pflueger, J., Lister, R., Rackham, O.J.L., Petretto, E., Polo, J.M., 2019. A single-cell atlas of entorhinal cortex from individuals with Alzheimer’s disease reveals cell-type-specific gene expression regulation. Nat. Neurosci. 22, 2087–2097. https://doi.org/10.1038/s41593-019-0539-4

Hardy, J., Allsop, D., 1991. Amyloid deposition as the central event in the aetiology of Alzheimer’s disease. Trends Pharmacol. Sci. 12, 383–388. https://doi.org/10.1016/0165-6147(91)90609-V

Hashimshony, T., Wagner, F., Sher, N., Yanai, I., 2012. CEL-Seq: Single-Cell RNA-Seq by Multiplexed Linear Amplification. Cell Rep. 2, 666–673. https://doi.org/10.1016/j.celrep.2012.08.003

Hemonnot, A.L., Hua, J., Ulmann, L., Hirbec, H., 2019. Microglia in Alzheimer disease: Well-known targets and new opportunities. Front. Aging Neurosci. 11, 1–20. https://doi.org/10.3389/fnagi.2019.00233

Hidalgo, J., Penkowa, M., Espejo, C., Martínez-Cáceres, E.M., Carrasco, J., Quintana, A., Molinero, A., Florit, S., Giralt, M., Ortega-Aznar, A., 2006. Expression of metallothionein-I,-II, and-III in Alzheimer disease and animal models of neuroinflammation. Exp. Biol. Med. 231, 1450–1458. https://doi.org/10.1177/153537020623100902

Hozumi, I., Asanuma, M., Yamada, M., Uchida, Y., 2004. Metallothioneins and neurodegenerative diseases. J. Heal. Sci. 50, 323–331. https://doi.org/10.1248/jhs.50.323

Hyman, B., Tanzi, R.E., 2019. Effects of Species-Specific Genetics on Alzheimer’s Mouse Models. Neuron 101, 351–352. https://doi.org/10.1016/j.neuron.2019.01.021

Itagaki, S., McGeer, P.L., Akiyama, H., Zhu, S., Selkoe, D., 1989. Relationship of microglia and astrocytes to amyloid deposits of Alzheimer disease. J. Neuroimmunol. 24, 173–182. https://doi.org/10.1016/0165-5728(89)90115-X

Jankowsky, J.L., Slunt, H.H., Ratovitski, T., Jenkins, N.A., Copeland, N.G., Borchelt, D.R., 2001. Co-expression of multiple transgenes in mouse CNS: A comparison of strategies. Biomol. Eng. 17, 157–165. https://doi.org/10.1016/S1389-0344(01)00067-3

Johnson, E.C.B., Dammer, E.B., Duong, D.M., Ping, L., Zhou, M., Yin, L., Higginbotham, L.A., Guajardo, A., White, B., Troncoso, J.C., Thambisetty, M., Montine, T.J., Lee, E.B., Trojanowski, J.Q., Beach, T.G., Reiman, E.M., Haroutunian, V., Wang, M., Schadt, E., Zhang, B., Dickson, D.W., Ertekin-Taner, N., Golde, T.E., Petyuk, V.A., De Jager, P.L., Bennett, D.A., Wingo, T.S., Rangaraju, S., Hajjar, I., Shulman, J.M., Lah, J.J., Levey, A.I., Seyfried, N.T., 2020. Large-scale proteomic analysis of Alzheimer’s disease brain and cerebrospinal fluid reveals early changes in energy metabolism associated with microglia and astrocyte activation. Nat. Med. 26. https://doi.org/10.1038/s41591-020-0815-6

Kamphuis, W., Kooijman, L., Orre, M., Stassen, O., Pekny, M., Hol, E.M., 2015. GFAP and Vimentin Deficiency Alters Gene Expression in Astrocytes and Microglia in Wild-Type Mice and Changes the Transcriptional Response of Reactive Glia in Mouse Model for Alzheimer’s Disease. Glia 63, 1036–1056. https://doi.org/10.1002/glia.22800

Kato, S., Gondo, T., Hoshii, Y., Takahashi, M., Yamada, M., Ishihara, T., 1998. Confocal observation of senile plaques in Alzheimer’s disease: Senile plaque morphology and relationship between senile plaques and astrocytes. Pathol. Int. 48, 332–340. https://doi.org/10.1111/j.1440-1827.1998.tb03915.x

Keren-Shaul, H., Spinrad, A., Weiner, A., Matcovitch-Natan, O., Dvir-Szternfeld, R., Ulland, T.K., David, E., Baruch, K., Lara-Astaiso, D., Toth, B., Itzkovitz, S., Colonna, M., Schwartz, M., Amit, I., 2017. A Unique Microglia Type Associated with Restricting Development of Alzheimer’s Disease. Cell 169, 1276–1290.e17. https://doi.org/10.1016/j.cell.2017.05.018

Kuleshov, M. V., Jones, M.R., Rouillard, A.D., Fernandez, N.F., Duan, Q., Wang, Z., Koplev, S., Jenkins, S.L., Jagodnik, K.M., Lachmann, A., McDermott, M.G., Monteiro, C.D., Gundersen, G.W., Ma’ayan, A., 2016. Enrichr: a comprehensive gene set enrichment analysis web server 2016 update. Nucleic Acids Res. 44, W90–W97. https://doi.org/10.1093/nar/gkw377

Lambert, J.C., Ibrahim-Verbaas, C.A., Harold, D., Naj, A.C., Sims, R., Bellenguez, C., Jun, G., DeStefano, A.L., Bis, J.C., Beecham, G.W., Grenier-Boley, B., Russo, G., Thornton-Wells, T.A., Jones, N., Smith, A. V., Chouraki, V., Thomas, C., Ikram, M.A., Zelenika, D., Vardarajan, B.N., Kamatani, Y., Lin, C.F., Gerrish, A., Schmidt, H., Kunkle, B., Fiévet, N., Amouyel, P., Pasquier, F., Deramecourt, V., De Bruijn, R.F.A.G., Amin, N., Hofman, A., Van Duijn, C.M., Dunstan, M.L., Hollingworth, P., Owen, M.J., O’Donovan, M.C., Jones, L., Holmans, P.A., Moskvina, V., Williams, J., Baldwin, C., Farrer, L.A., Choi, S.H., Lunetta, K.L., Fitzpatrick, A.L., Harris, T.B., Psaty, B.M., Gilbert, J.R., Hamilton-Nelson, K.L., Martin, E.R., Pericak-Vance, M.A., Haines, J.L., Gudnason, V., Jonsson, P. V., Eiriksdottir, G., Bihoreau, M.T., Lathrop, M., Valladares, O., Cantwell, L.B., Wang, L.S., Schellenberg, G.D., Ruiz, A., Boada, M., Reitz, C., Mayeux, R., Ramirez, A., Maier, W., Hanon, O., Kukull, W.A., Buxbaum, J.D., Campion, D., Wallon, D., Hannequin, D., Crane, P.K., Larson, E.B., Becker, T., Cruchaga, C., Goate, A.M., Craig, D., Johnston, J.A., McGuinness, B., Todd, S., Passmore, P., Berr, C., Ritchie, K., Lopez, O.L., De Jager, P.L., Evans, D., Lovestone, S., Proitsi, P., Powell, J.F., Letenneur, L., Barberger-Gateau, P., Dufouil, C., Dartigues, J.F., Morón, F.J., Rubinsztein, D.C., St. George-Hyslop, P., Sleegers, K., Bettens, K., Van Broeckhoven, C., Huentelman, M.J., Gill, M., Brown, K., Morgan, K., Kamboh, M.I., Keller, L., Fratiglioni, L., Green, R., Myers, A.J., Love, S., Rogaeva, E., Gallacher, J., Bayer, A., Clarimon, J., Lleo, A., Tsuang, D.W., Yu, L., Bennett, D.A., Tsolaki, M., Bossù, P., Spalletta, G., Collinge, J., Mead, S., Sorbi, S., Nacmias, B., Sanchez-Garcia, F., Deniz Naranjo, M.C., Fox, N.C., Hardy, J., Bosco, P., Clarke, R., Brayne, C., Galimberti, D., Mancuso, M., Matthews, F., Moebus, S., Mecocci, P., Del Zompo, M., Hampel, H., Pilotto, A., Bullido, M., Panza, F., Caffarra, P., Mayhaus, M., Pichler, S., Gu, W., Riemenschneider, M., Lannfelt, L., Ingelsson, M., Hakonarson, H., Carrasquillo, M.M., Zou, F., Younkin, S.G., Beekly, D., Alvarez, V., Coto, E., Razquin, C., Pastor, P., Mateo, I., Combarros, O., Faber, K.M., Foroud, T.M., Soininen, H., Hiltunen, M., Blacker, D., Mosley, T.H., Graff, C., Holmes, C., Montine, T.J., Rotter, J.I., Brice, A., Nalls, M.A., Kauwe, J.S.K., Boerwinkle, E., Schmidt, R., Rujescu, D., Tzourio, C., Nöthen, M.M., Launer, L.J., Seshadri, S., 2013. Meta-analysis of 74,046 individuals identifies 11 new susceptibility loci for Alzheimer’s disease. Nat. Genet. 45, 1452–1458. https://doi.org/10.1038/ng.2802

Leone, C., Le Pavec, G., Meme, W., Porcheray, F., Samah, B., Dormont, D., Gras, G., 2006. Characterization of human monocyte-derived microglia-like cells. Glia 54, 183–192. https://doi.org/10.1002/glia

Li, H., Durbin, R., 2010. Fast and accurate long-read alignment with Burrows-Wheeler transform. Bioinformatics 26, 589–595. https://doi.org/10.1093/bioinformatics/btp698

Lue, L.F., Beach, T.G., Walker, D.G., 2019. Alzheimer’s Disease Research Using Human Microglia. Cells 8, 1– 19. https://doi.org/10.3390/cells8080838

Manso, Y., Carrasco, J., Comes, G., Adlard, P.A., Bush, A.I., Hidalgo, J., 2012. Characterization of the role of the antioxidant proteins metallothioneins 1 and 2 in an animal model of Alzheimer’s disease. Cell. Mol. Life Sci. 69, 3665–3681. https://doi.org/10.1007/s00018-012-1045-y

Massaad, C.A., Washington, T.M., Pautler, R.G., Klann, E., 2009. Overexpression of SOD-2 reduces hippocampal superoxide and prevents memory deficits in a mouse model of Alzheimer’s disease. Proc. Natl. Acad. Sci. U. S. A. 106, 13576–13581. https://doi.org/10.1073/pnas.0902714106

Mathys, H., Davila-Velderrain, J., Peng, Z., Gao, F., Mohammadi, S., Young, J.Z., Menon, M., He, L., Abdurrob, F., Jiang, X., Martorell, A.J., Ransohoff, R.M., Hafler, B.P., Bennett, D.A., Kellis, M., Tsai, L.H., 2019. Single-cell transcriptomic analysis of Alzheimer’s disease. Nature 570, 332–337. https://doi.org/10.1038/s41586-019-1195-2

Melief, J., Sneeboer, M.A.M., Litjens, M., Ormel, P.R., Palmen, S.J.M.C., Huitinga, I., Kahn, R.S., Hol, E.M., de Witte, L.D., 2016. Characterizing primary human microglia: A comparative study with myeloid subsets and culture models. Glia 64, 1857–1868. https://doi.org/10.1002/glia.23023

Muraro, M.J., Dharmadhikari, G., Grün, D., Groen, N., Dielen, T., Jansen, E., van Gurp, L., Engelse, M.A., Carlotti, F., de Koning, E.J.P., van Oudenaarden, A., 2016. A Single-Cell Transcriptome Atlas of the Human Pancreas. Cell Syst. 3, 385–394.e3. https://doi.org/10.1016/j.cels.2016.09.002

Nunomura, A., Castellani, R.J., Zhu, X., Moreira, P.I., Perry, G., Smith, M.A., 2006. Involvement of oxidative stress in Alzheimer disease. J. Neuropathol. Exp. Neurol. 65, 631–641. https://doi.org/10.1097/01.jnen.0000228136.58062.bf

Ohgidani, M., Kato, T.A., Kanba, S., 2015. Introducing directly induced microglia-like (iMG) cells from fresh human monocytes: A novel translational research tool for psychiatric disorders. Front. Cell. Neurosci. 9, 1– 5. https://doi.org/10.3389/fncel.2015.00184

Ormel, P.R., Böttcher, C., Gigase, F.A.J., Missall, R.D., van Zuiden, W., Fernández Zapata, M.C., Ilhan, D., de Goeij, M., Udine, E., Sommer, I.E.C., Priller, J., Raj, T., Kahn, R.S., Hol, E.M., de Witte, L.D., 2020. A characterization of the molecular phenotype and inflammatory response of schizophrenia patient-derived microglia-like cells. Brain. Behav. Immun. 90, 196–207. https://doi.org/10.1016/j.bbi.2020.08.012

Orre, M., Kamphuis, W., Osborn, L.M., Jansen, A.H.P.P., Kooijman, L., Bossers, K., Hol, E.M., 2014. Isolation of glia from Alzheimer’s mice reveals inflammation and dysfunction. Neurobiol. Aging 35, 2746–2760. https://doi.org/10.1016/j.neurobiolaging.2014.06.004

Pedersen, M., Jensen, R., Pedersen, D.S., Skjolding, A.D., Hempel, C., Maretty, L., Penkowa, M., 2009. Metallothionein-I+II in neuroprotection. BioFactors 35, 315–325. https://doi.org/10.1002/biof.44

Prokop, S., Miller, K.R., Heppner, F.L., 2013. Microglia actions in Alzheimer’s disease. Acta Neuropathol. 126, 461–477. https://doi.org/10.1007/s00401-013-1182-x

Querfurth, H.W., LaFerla, F.M., 2010. Alzheimer’s disease. N. Engl. J. Med. 364, 329–344. https://doi.org/10.1007/978-1-4939-7880-9_9

Raina, A.K., Templeton, D.J., Deak, J.C., Perry, G., Smith, M.A., 1999. Quinone reductase (NQO1), a sensitive redox indicator, is increased in Alzheimer’s disease. Redox Rep. 4, 23–27. https://doi.org/10.1179/135100099101534701

Rogers, J., Strohmeyer, R., Kovelowski, C.J., Li, R., 2002. Microglia and inflammatory mechanisms in the clearance of amyloid β peptide. Glia 40, 260–269. https://doi.org/10.1002/glia.10153

Rozemuller, J.M., Eikelenboom, P., Stam, F.C., 1986. Role of microglia in plaque formation in senile dementia of the Alzheimer type - An immunohistochemical study. Virchows Arch. B Cell Pathol. Incl. Mol. Pathol. 51, 247–254. https://doi.org/10.1007/BF02899034

Ryan, K.J., White, C.C., Patel, K., Xu, J., Olah, M., Replogle, J.M., Frangieh, M., Cimpean, M., Winn, P., McHenry, A., Kaskow, B.J., Chan, G., Cuerdon, N., Bennett, D.A., Boyd, J.D., Imitola, J., Elyaman, W., De Jager, P.L., Bradshaw, E.M., 2017. A human microglia-like cellular model for assessing the effects of neurodegenerative disease gene variants. Sci. Transl. Med. 9, 1–13. https://doi.org/10.1126/scitranslmed.aai7635

Salminen, A., Ojala, J., Kauppinen, A., Kaarniranta, K., Suuronen, T., 2009. Inflammation in Alzheimer’s disease: Amyloid-β oligomers trigger innate immunity defence via pattern recognition receptors. Prog. Neurobiol. 87, 181–194. https://doi.org/10.1016/j.pneurobio.2009.01.001

SantaCruz, K.S., Yazlovitskaya, E., Collins, J., Johnson, J., DeCarli, C., 2004. Regional NAD(P)H:quinone oxidoreductase activity in Alzheimer’s disease. Neurobiol. Aging 25, 63–69. https://doi.org/10.1016/S0197-4580(03)00117-9

Sarlus, H., Heneka, M.T., 2017. Microglia in Alzheimer ‘ s disease. J. Clin. Invest. 127, 3240–3249. https://doi.org/10.1172/JCI90606

Searle, P.F., Davison, B.L., Stuart, G.W., Wilkie, T.M., Norstedt, G., Palmiter, R.D., 1984. Regulation, linkage, and sequence of mouse metallothionein I and II genes. Mol. Cell. Biol. 4, 1221–1230. https://doi.org/10.1128/mcb.4.7.1221

Selkoe, D.J., 1991. The Molecular Pathology of Alzheimer’s Disease. Neuron 6, 487–498. https://doi.org/10.1016/0896-6273(91)90052-2

Selkoe, D.J., Hardy, J., 2016. The amyloid hypothesis of Alzheimer’s disease at 25 years. EMBO Mol. Med. 8, 595–608. https://doi.org/10.15252/emmm.201606210

Sellgren, C.M., Gracias, J., Watmuff, B., Biag, J.D., Thanos, J.M., Whittredge, P.B., Fu, T., Worringer, K., Brown, H.E., Wang, J., Kaykas, A., Karmacharya, R., Goold, C.P., Sheridan, S.D., Perlis, R.H., 2019. Increased synapse elimination by microglia in schizophrenia patient-derived models of synaptic pruning. Nat. Neurosci. https://doi.org/10.1038/s41593-018-0334-7

Simmini, S., Bialecka, M., Huch, M., Kester, L., Van De Wetering, M., Sato, T., Beck, F., Van Oudenaarden, A., Clevers, H., Deschamps, J., 2014. Transformation of intestinal stem cells into gastric stem cells on loss of transcription factor Cdx2. Nat. Commun. 5, 1–10. https://doi.org/10.1038/ncomms6728

Sneeboer, M.A.M., Snijders, G.J.L.J., Berdowski, W.M., Fernández-Andreu, A., van Mierlo, H.C., Berdenis van Berlekom, A., Litjens, M., Kahn, R.S., Hol, E.M., de Witte, L.D., 2019. Microglia in post-mortem brain tissue of patients with bipolar disorder are not immune activated. Transl. Psychiatry 9. https://doi.org/10.1038/s41398-019-0490-x

Song, M., Jin, J., Lim, J.E., Kou, J., Pattanayak, A., Rehman, J.A., Kim, H.D., Tahara, K., Lalonde, R., Fukuchi, K., 2011. TLR4 mutation reduces microglial activation, increases Abeta deposits and exacerbates cognitive deficits in a mouse model of Alzheimer’s disease. J. Neuroinflammation 8, 1–14. https://doi.org/10.1186/1742-2094-8-92

Srinivasan, K., Friedman, B.A., Etxeberria, A., Huntley, M.A., Brug M.P. van der, Foreman, O., Paw, J.S., Modrusan, Z., Beach, T., Serrano, G., Hansen, D., 2019. Alzheimer’s patient brain myeloid cells exhibit enhanced aging and unique transcriptional activation. bioRxiv 610345. https://doi.org/10.1101/610345

Tahara, K., Kim, H.D., Jin, J.J., Maxwell, J.A., Li, L., Fukuchi, K., 2006. Role of toll-like receptor signalling in Aβ uptake and clearance. Brain 129, 3006–3019. https://doi.org/10.1093/brain/awl249

Tejera, D., Heneka, M.T., 2016. Microglia in Alzheimer’s Disease: The Good, the Bad and the Ugly. Curr. Alzheimer Res. 13, 370–380. https://doi.org/10.2174/1567205013666151116125012

Van Tijn, P., Dennissen, F.J.A.A., Gentier, R.J.G.G., Hobo, B., Hermes, D., Steinbusch, H.W.M., Van Leeuwen, F.W., Fischer, D.F., 2012. Mutant ubiquitin decreases amyloid β plaque formation in a transgenic mouse model of Alzheimer’s disease. Neurochem. Int. 61, 739–748. https://doi.org/10.1016/j.neuint.2012.07.007

Vašák, M., Meloni, G., 2011. Chemistry and biology of mammalian metallothioneins. J. Biol. Inorg. Chem. 16, 1067–1078. https://doi.org/10.1007/s00775-011-0799-2

Walker, D.G., Link, J., Lue, L.-F., Dalsing-Hernandez, J.E., Boyes, B.E., 2006. Gene expression changes by amyloid β peptide-stimulated human postmortem brain microglia identify activation of multiple inflammatory processes. J. Leukoc. Biol. 79, 596–610. https://doi.org/10.1189/jlb.0705377

Walker, D.G., Lue, L.F., Beach, T.G., 2001. Gene expression profiling of amyloid beta peptide-stimulated human post-mortem brain microglia. Neurobiol. Aging 22, 957–966. https://doi.org/10.1016/S0197-4580(01)00306-2

Waller, R., Murphy, M., Garwood, C.J., Jennings, L., Heath, P.R., Chambers, A., Matthews, F.E., Brayne, C., Ince, P.G., Wharton, S.B., Simpson, J.E., 2018. Metallothionein-I/II expression associates with the astrocyte DNA damage response and not Alzheimer-type pathology in the aging brain. Glia 66, 2316–2323. https://doi.org/10.1002/glia.23465

West, A.K., Hidalgo, J., Eddins, D., Levin, E.D., Aschner, M., 2008. Metallothionein in the central nervous system: Roles in protection, regeneration and cognition. Neurotoxicology 29, 489–503. https://doi.org/10.1016/j.neuro.2007.12.006

Wolf, S.A., Boddeke, H.W.G.M., Kettenmann, H., 2017. Microglia in Physiology and Disease. Annu. Rev. Physiol. 79, 619–643. https://doi.org/10.1146/annurev-physiol-022516-034406

Zambenedetti, P., Giordano, R., Zatta, P., 1998. Metallothioneins are highly expressed in astrocytes and microcapillaries in Alzheimer’s disease. J. Chem. Neuroanat. 15, 21–26. https://doi.org/10.1016/S0891-0618(98)00024-6

